# Fetal and trophoblast PI3Kp110α have distinct roles in regulating resource supply to the growing fetus

**DOI:** 10.1101/473967

**Authors:** Jorge Lopez-Tello, Vicente Perez-Garcia, Jaspreet Khaira, Laura C. Kusinski, Wendy N. Cooper, Adam Andreani, Imogen Grant, Edurne Fernandez de Liger, Myriam Hemberger, Ionel Sandovici, Miguel Constancia, Amanda N. Sferruzzi-Perri

**Affiliations:** Centre for Trophoblast Research, Department of Physiology, Development and Neuroscience, Downing Street, University of Cambridge, Cambridge, UK CB2 3EG; Epigenetics Programme, The Babraham Institute, Babraham Research Campus, Cambridge CB22 3AT, UK; Metabolic Research Laboratories, MRC Metabolic Diseases Unit, Department of Obstetrics and Gynaecology, The Rosie Hospital, Robinson Way, Cambridge, UK CB2 0SW; Departments of Biochemistry & Molecular Biology and Medical Genetics, Cumming School of Medicine, University of Calgary, Calgary T2N 4N1, Alberta, Canada

## Abstract

Previous studies suggest that the placental supply of nutrients to the fetus adapts according to fetal demand. However, the signaling events underlying placental adaptations remain largely unknown. Earlier work in mice has revealed that loss of the phosphoinositide 3-kinase p110α impairs feto-placental growth but placental nutrient supply is adaptively increased. Here we explore the role of p110α in the epiblast-derived (fetal) and trophoblast lineages of the conceptus in relation to feto-placental growth and placental development and transfer function. Using conditional gene manipulations to knock-down p110α either by ∼50% or ∼100% in the fetal lineages and/or trophoblast, this study shows that p110α in the fetus is essential for prenatal development and a major regulator of placental phenotype in mice. Complete loss of fetal p110α caused embryonic death, whilst heterozygous loss resulted in fetal growth restriction and impaired placental formation and nutrient transport. Loss of trophoblast p110α also resulted in abnormal placental development, although fetuses were viable. However, in response to complete loss of trophoblast p110α, the placenta failed to transport sufficient amino acid to match fetal demands for growth. Using RNA-seq, we identified several genes downstream of p110α in the trophoblast that are important in adapting placental phenotype to support fetal growth. Further work using CRISPR/Cas9 genome targeting showed that loss of p110α differentially affects the expression of genes in trophoblast and embryonic stem cells. Our findings thus reveal important, but distinct roles for p110α signaling in the different compartments of the conceptus, which control fetal resource acquisition and ultimately affect healthy growth.

**One Sentence Summary:** Fetal and trophoblast p110α modify resource allocation

## Introduction

Intrauterine growth is dictated by the genetically-determined fetal drive for growth and the supply of nutrients and oxygen to the fetus. In turn, fetal substrate supply depends on the functional capacity of the placenta to transfer nutrients and oxygen from the mother to the fetus. Insufficient fetal substrate supply prevents the fetus from achieving its genetic growth potential and leads to intrauterine growth restriction, which affects up to 10% of the population and is associated with perinatal morbidity and mortality (Baschat et al., 2007; Baschat and Hecher, 2004). Studies have shown that the transport capacity of the placenta is diminished in human pregnancies associated with fetal growth restriction (Glazier et al., 1997; Jansson and Powell, 2006), suggesting that placental insufficiency may be the underlying cause of a fetus failing to achieve its genetic growth potential. However, there is also evidence that placental transport capacity may adapt to maintain the supply of resources appropriate for the growth potential of the fetus (Sandovici et al., 2012; Sferruzzi-Perri and Camm, 2016). For instance, the placental capacity to supply resources to the fetus is higher in the lightest compared to heaviest placentas supporting babies within the normal birth weight range (Godfrey et al., 1998) and is increased in newborns showing some protection to the growth-restricting effects of altitude-induced hypoxia (Espinoza et al., 2001; Jackson et al., 1988; Moore et al., 2001). Studies in mice have also shown that the placenta can adaptively up-regulate its transport function when the fetal demand for resources exceeds the supply capacity of the placenta, including following deletion of nutrient transporter or growth-regulatory genes, in naturally small placentas within litters, and when the maternal provision of substrates is suboptimal (Coan et al., 2008; Constancia et al., 2005; Dilworth et al., 2010; Higgins et al., 2015; Sferruzzi-Perri et al., 2011; Sferruzzi-Perri et al., 2013b; Vaughan et al., 2013; Wyrwoll et al., 2009). Taken together, these observations suggest that the placenta may adopt compensatory mechanisms to help meet the fetal genetic drive for growth, and that perhaps in severe cases of fetal growth restriction such adaptive attempts fail to occur or are insufficient. However, the signaling events which underlie placental adaptations require elucidation.

The phosphoinositide 3-kinases (PI3Ks) are a highly conserved family of enzymes, which are thought to have evolved to regulate growth in relation to nutrient supply (Engelman et al., 2006; Kriplani et al., 2015). In response to receptor activation, they generate lipid second messengers to initiate signaling cascades, which control important physiological processes as diverse as cellular proliferation, metabolism, survival, polarity and membrane trafficking (Vanhaesebroeck et al., 2010). They are grouped into three PI3K classes (I, II and III) on the basis of their substrate preference and sequence similarity and in mammals, Class I PI3Ks can be further subdivided according to the receptors to which they couple. Class IA PI3Ks have received the most attention to date and signal downstream of growth factor receptor tyrosine kinases (Jean and Kiger, 2014). In adult mammalian tissues, the ubiquitously expressed Class IA isoform, p110α plays a key role in mediating the growth and metabolic effects of insulin and insulin-like growth factors (IGFs) (Foukas et al., 2006; Knight et al., 2006; Sopasakis et al., 2010), which are major growth factors that also operate during feto-placental development (Sferruzzi-Perri et al., 2017; Sferruzzi-Perri et al., 2013a). Work in mice has shown that changes in the PI3K pathway, particularly downstream of p110α were observed in placentas showing adaptive up-regulation of substrate supply to the fetus in pregnancies challenged by poor maternal environments or genetically-induced placental growth restriction (Higgins et al., 2015; Sferruzzi-Perri et al., 2011; Sferruzzi-Perri et al., 2013b). These observations indicate that p110α may be part of a critical signaling cascade that integrates the various environmental signals involved in adapting placental resource allocation to fetal growth.

Studies in mice have indicated that embryos die between day 9.5 and 11.5 of pregnancy when they are homozygous deficient for p110α by either deleting the *Pik3ca* gene or by a mutation in *Pik3ca*, rendering p110α inactive (kinase dead mutation; p110α^D933A/D933A^; α/α) (Bi et al., 1999; Foukas et al., 2006; Graupera et al., 2008). When p110α activity is reduced by half, in mice carrying one copy of the kinase-dead mutation (p110α^D933A/+^; α/+), fetuses are viable but weigh 85-90% of wildtype in late gestation (Foukas et al., 2006; Sferruzzi-Perri et al., 2016). Placental weight is also reduced in association with defects in the development of the labyrinthine transport region in α/+ mutants (Sferruzzi-Perri et al., 2016). However, the capacity of the placenta to transfer maternal glucose and amino acid (via System A) to the fetus per unit surface area is increased in α/+ mutants near term (Sferruzzi-Perri et al., 2016). This suggests adaptation of placental supply capacity with p110α deficiency. In the α/+ mutant the whole conceptus is heterozygous for the p110α mutation (Sferruzzi-Perri et al., 2016) and the contribution of the fetal *versus* the trophoblast cell lineages in driving changes in placental development and adaptive responses cannot be discerned. This study therefore, sought to answer the following questions: 1) what effect does fetal *versus* trophoblast p110α deficiency have on placental phenotype and resource allocation to fetal growth? 2) does adaptive upregulation of placental transport depend on p110α in the fetal or trophoblast lineages? and 3) which genes downstream of p110α in the trophoblast compartment of the conceptus are implicated in adapting placental phenotype to support fetal growth in late mouse pregnancy? To achieve this, using conditional deletions, we selectively manipulated the expression of the p110α gene (*Pik3ca*) in the trophoblast and/or fetal cell lineages and assessed the impact on placental morphology, transport and transcriptome in relation to fetal growth.

## Results

### Fetal and trophoblast p110α interplay to regulate growth of the conceptus

We found PI3K p110α was highly expressed by the placental transport labyrinthine zone and the endocrine junctional zone of the mouse placenta (Fig. S1). We then halved p110α expression in either the trophoblast or the fetal (epiblast-derived) lineages of the conceptus by mating mice in which exons 18-19 of the *Pik3ca* gene were flanked by LoxP sites (Graupera et al., 2008), to *Cyp19*Cre (Wenzel and Leone, 2007) and *Meox2*Cre (Tallquist and Soriano, 2000) mice, respectively, termed throughout as Het-P and Het-F. These Cre lines are active in opposite compartments of the conceptus; for Het-P the *Cyp19*Cre targets expression in the trophoblast lineages of the placenta but not fetus, labyrinthine fetal capillaries or mesenchyme in the chorion (Fig. 1A and C). In contrast, in Het-F the *Meox2*Cre deletes p110α expression in the fetus and placental labyrinthine fetal capillaries and chorionic mesenchyme but not trophoblast (Fig. 1B and C) (further information regarding the genetic crosses can be found in Table S1). We then compared the fetal and placental growth phenotype of the conditional Het-P and Het-F on day 19 of pregnancy (term ∼20 days) to conceptuses with global heterozygous p110α deficiency, achieved with the ubiquitous *CMV*Cre line (Schwenk et al., 1995) (termed Het-U, Fig. 1C). We found that compared to their wild-type (WT) littermates, there was no effect of heterozygous deficiency of p110α in the trophoblast on fetal or placental weight in Het-P mutants (Fig. 2A). However, fetal and placental weights were 10-15% lighter for Het-F and Het-U conceptuses (Fig. 2B and C). The findings in Het-F and Het-U mutants are consistent with the proliferation defects observed in embryos with a deficiency in p110α (Bi et al., 1999; Foukas et al., 2006) and reveal for the first time that p110α in the embryo is important for determining the size of the placenta.

**Fig. 1.**
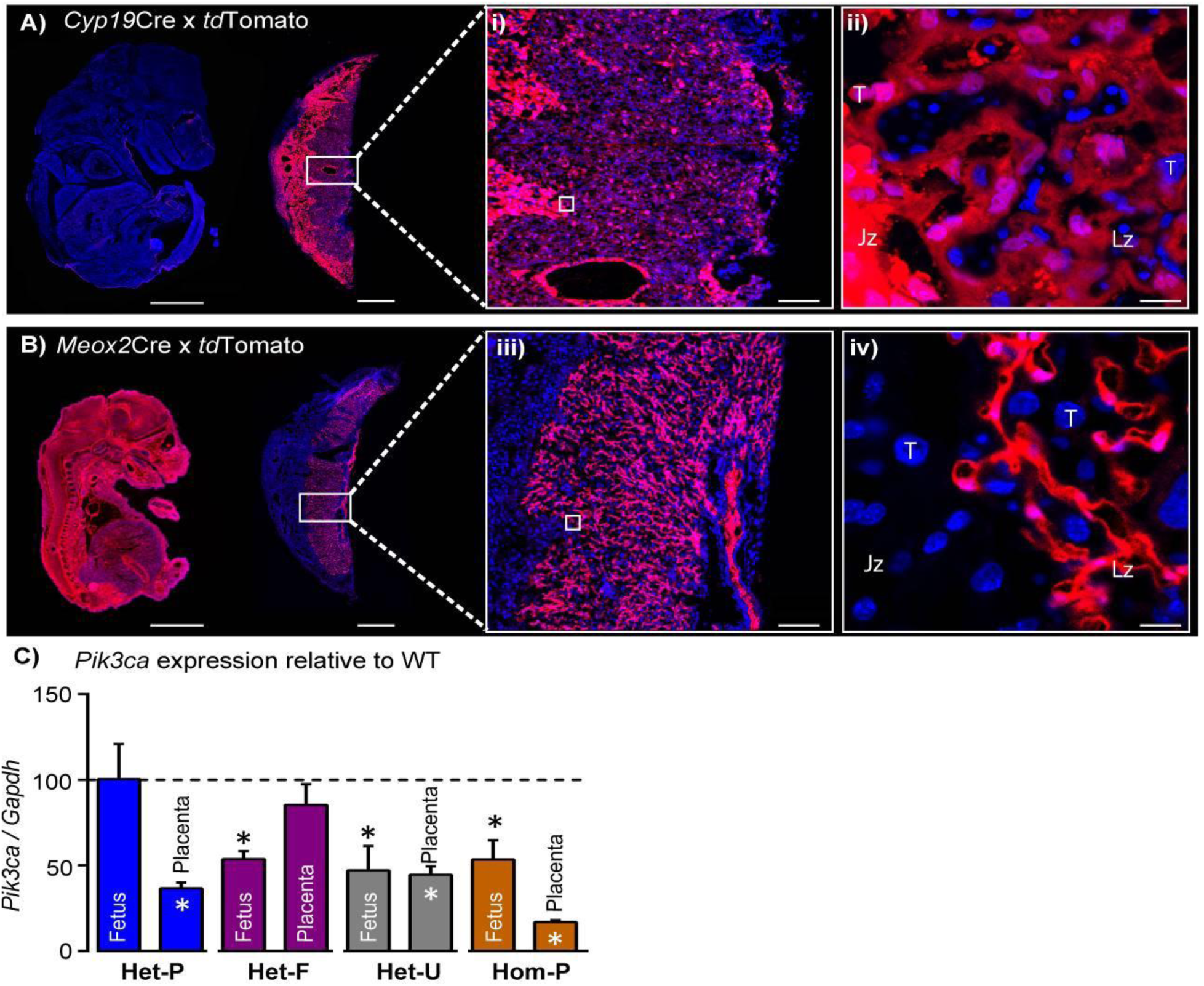
Successful modulation of p110α expression in the conceptus. (**A-B**) Validation that C*yp19*Cre (**A**) and *Meox2*Cre (**B**) are active in opposite compartments in the conceptus by crossing lines to the *tdTomato* reporter and assessing placentas and fetuses on day 16 of pregnancy. Boxes in A and B are shown in high magnification in i and ii, respectively. Scale bar for fetuses and placentas in A and B = 2mm and 1mm, in i and ii = 200μm, and in iii and iv = 20μm, respectively. (**C**) qRT-PCR for *Pik3ca* gene normalised to *Gapdh* in whole homogenates of fetus and placenta of Het-P, Het-F, Het-U and Hom-P mutants on day 19 of pregnancy, expressed as a ratio of their respective wild-type control (WT, denoted as a dotted line). *Gapdh* expression was not affected by genotype. *P < 0.05, **P < 0.01, and ***P < 0.001 *versus* WT, unpaired t test. n≥4 per genotype. Het-F=heterozygous deficiency in the fetus, Het-P=heterozygous deficiency in the placenta, Het-U=heterozygous deficiency in the fetus and placenta, Hom-P=heterozygous deficiency in the fetus and homozygous deficiency in the placenta, Jz=Junctional zone, Lz=Labyrinth zone, T=Trophoblast.

**Fig. 2.**
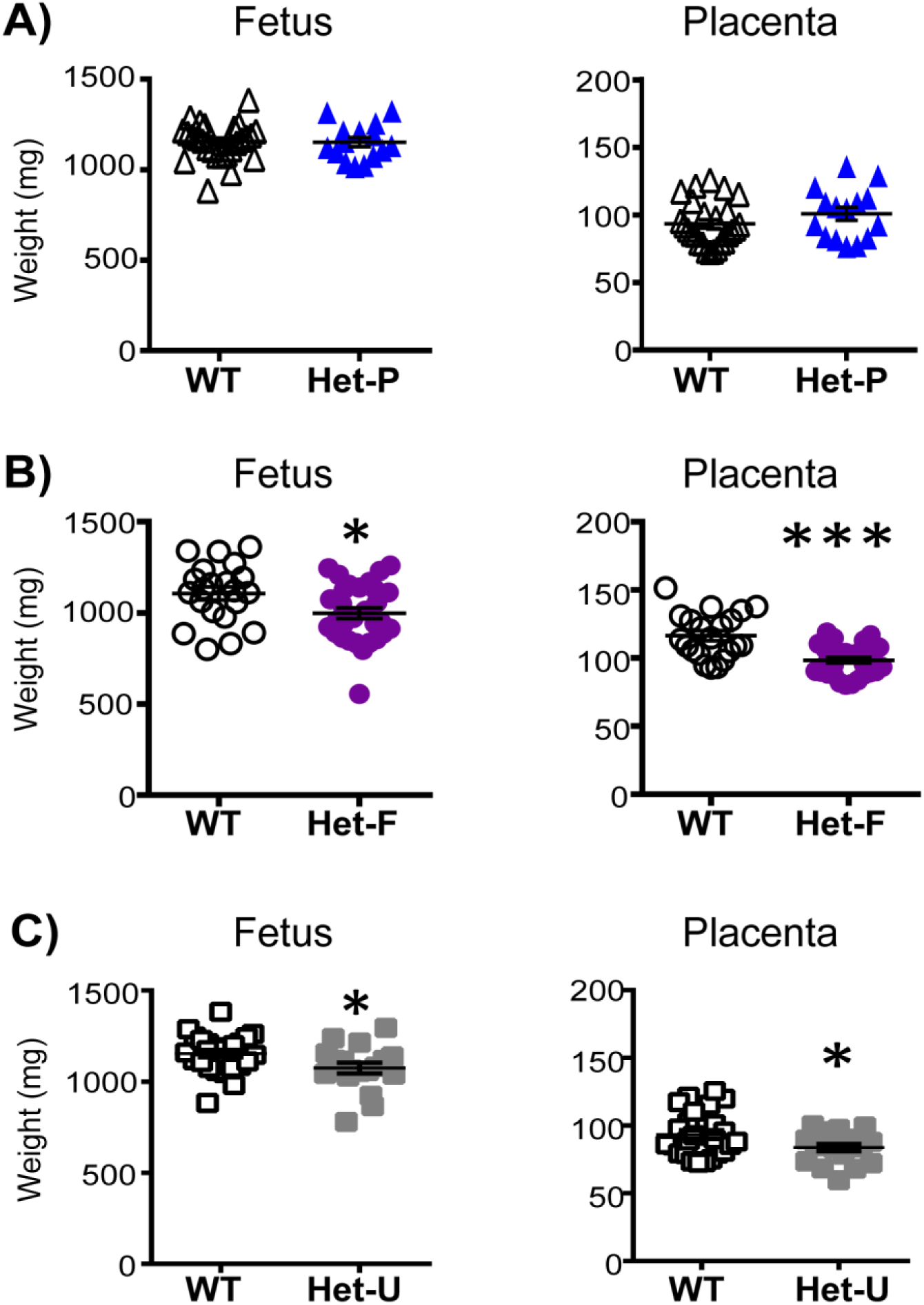
Feto-placental growth is regulated by fetal but not trophoblast p110α. (**A**) Het-P (n=15) compared to WT (n=26) (**B**), Het-F (n=30) compared to WT (n=21) and (**C**) Het-U (n=18) compared to WT (n=26) on day 19 of pregnancy. *P < 0.05 and ***P < 0.001 *versus* WT, unpaired t test. Note, there was no significant difference in fetal weight between Het-F and WT when data were analyzed by paired t test. Data from individual conceptuses are shown and the mean is denoted as a horizontal line with SEM.

Although fetal and not trophoblast p110α deficiency reduced conceptus weight, both affected the structure of the placenta (Fig. 3A-C). In the labyrinthine region of Het-P conceptuses, the volume of maternal blood spaces, fetal capillaries and surface area were reduced though trophoblast increased compared to their WT controls (Fig. 3A). In Het-F conceptuses, the labyrinthine zone, fetal capillaries and trophoblast volume were decreased *versus* WT littermates (Fig. 3B). In the ubiquitous p110α heterozygote (Het-U), the volume of the labyrinthine zone, fetal capillaries and trophoblast were reduced, surface area decreased and barrier to diffusion was greater, relative to WT littermates (Fig. 3C). Reassuringly, the same defects in Het-U placental structure were previously observed for α/+ mutants near term (Sferruzzi-Perri et al., 2016). The volume of the endocrine junctional zone in Het-P, Het-F or Het-U placentas was not significantly altered when compared to the respective WT controls (Fig. S2). Taken together, these findings indicate that p110α in the fetal and trophoblast lineages of the conceptus interplay to regulate the development of the transport region in the placenta.

**Fig. 3.**
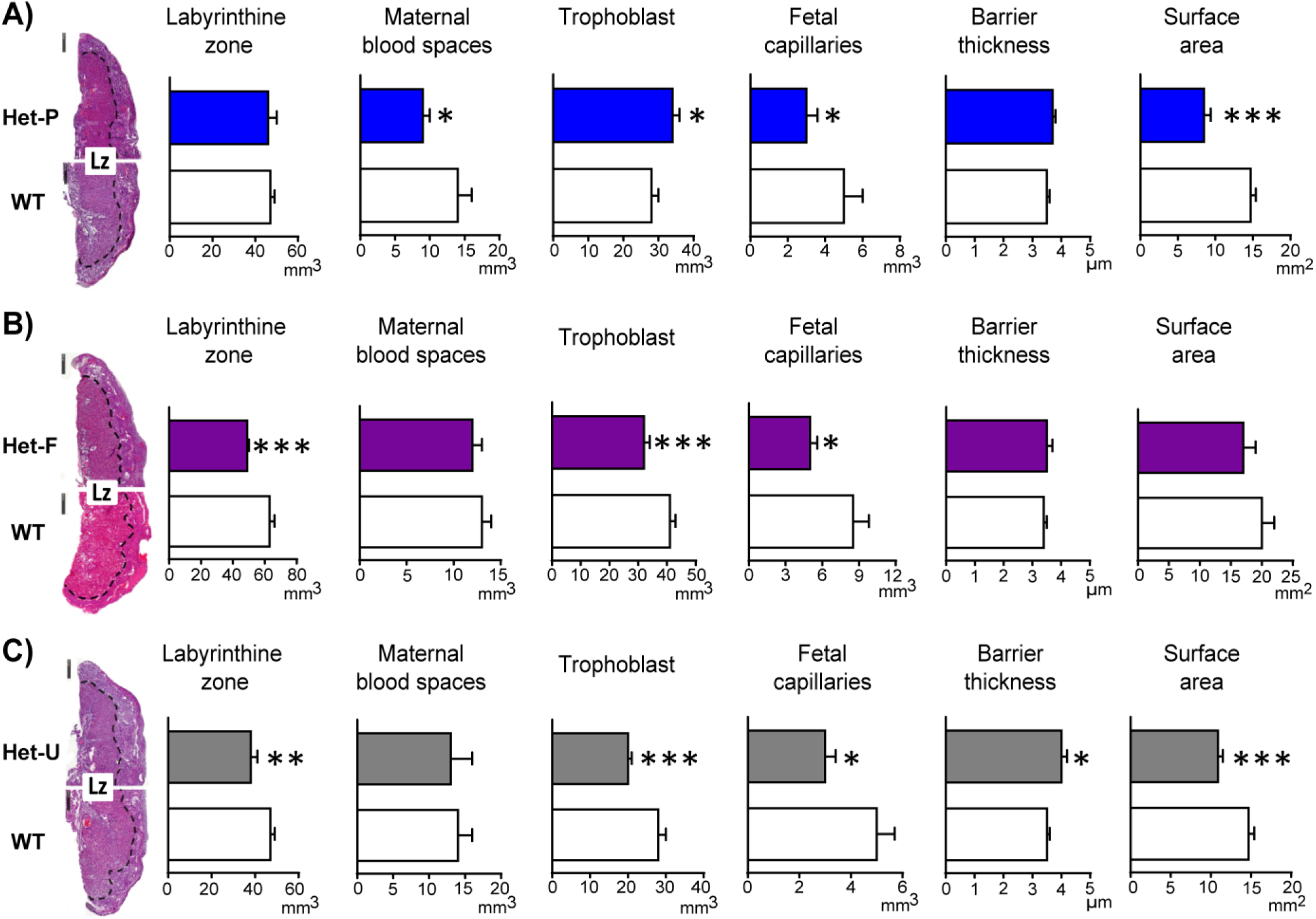
Placental morphology is regulated by both fetal and trophoblast p110α. (**A**) Het-P, (**B**) Het-F and (**C**) Het-U placental morphology on day 19 of pregnancy as determined by stereology. *P < 0.05, **P < 0.01, and ***P < 0.001 *versus* WT, unpaired t test. Data presented as means ± SEM. Scale bar for hematoxylin and eosin-stained placental cross-sections is 500μm. 6-10 placentas were assessed for each genotype for labyrinthine zone (Lz) volume and 4-5 analysed for Lz morphology.

### Fetal and trophoblast p110α both regulate placental resource allocation to fetal growth

To assess whether morphological alterations in the placenta affect placental resource allocation to the fetus in Het-F, Het-U and Het-P, we measured the uni-directional maternal-fetal transfer of the non-metabolisable analogues of glucose (^3^H-methyl-D glucose; MeG) and a neutral amino acid (^14^C-methyl amino-isobutyric acid; MeAIB) on day 19 of pregnancy. We assessed fetal counts in relation to the estimated surface area for transport or to fetal weight, which respectively provided us with indices of the placental capacity for nutrient transfer and fetal growth relative to supply. We found that in compensation for the impaired labyrinthine development, Het-P and Het-U placentas transferred more MeG and MeAIB and Het-F more MeAIB per unit surface area compared to their respective WT littermates (Fig. 4A-C). In agreement with these findings, mutant fetuses received either normal (MeG in all mutants and MeAIB in Het-P and Het-F) or increased amount of solutes (MeAIB in Het-U) for their size (Fig. 4D-F). Moreover, the total fetal accumulation of these solutes was the same in the mutant and WT fetuses (Fig. S3). These data suggest an adaptive response of both facilitated (MeG) and active (MeAIB) transport functions in placentas that are morphologically compromised by a lack of fetal or trophoblast p110α.

**Fig. 4.**
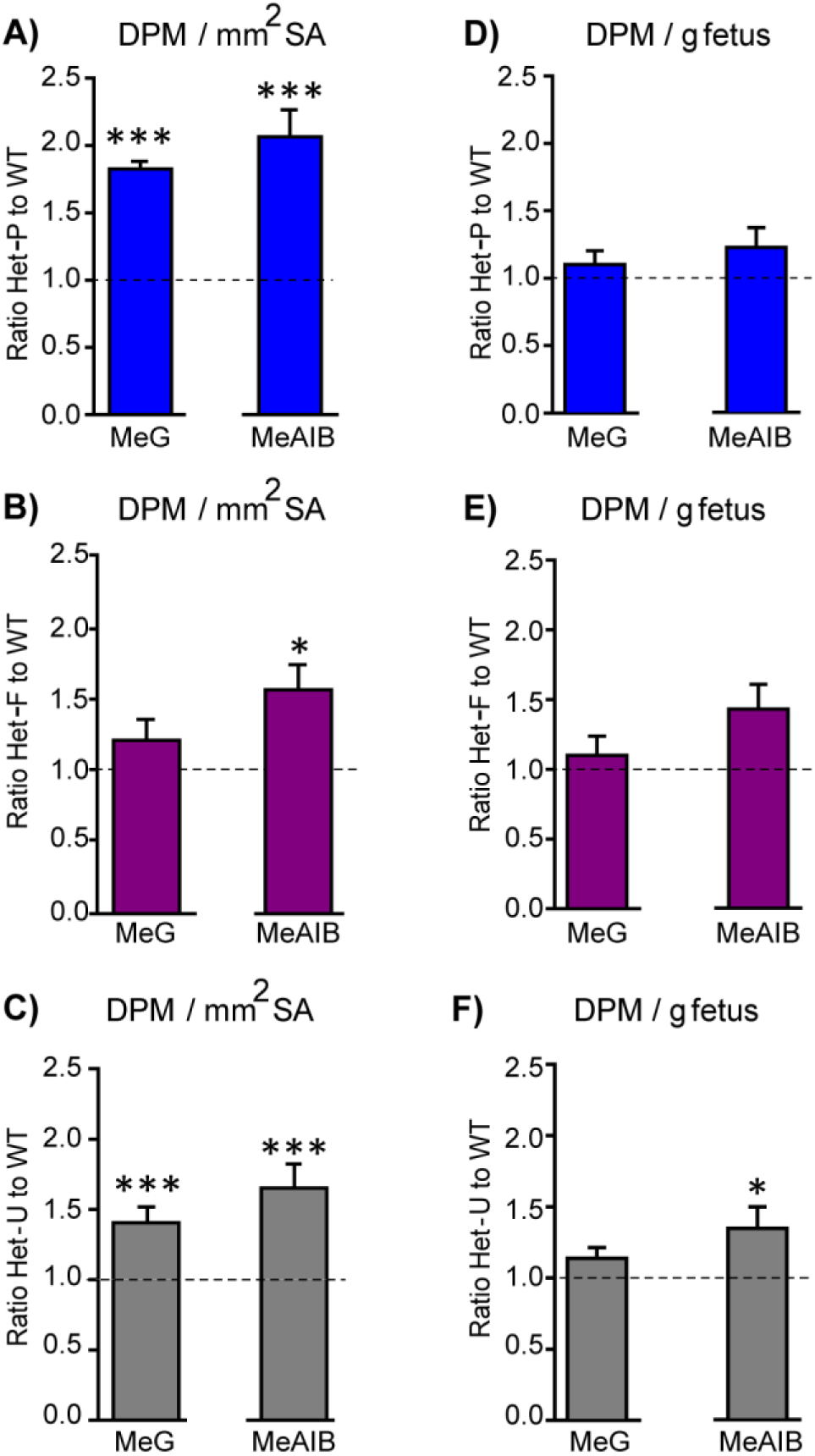
Placental nutrient transfer is adaptively increased in response to fetal and/or trophoblast loss of p110α. (**A-F**) The capacity of the placenta to transport ^3^H-methyl-D glucose (MeG) and ^14^C-amino isobutyric acid (MeAIB) relative to surface area available or to fetal weight (**D-F**) on day 19 of pregnancy for Het-P (n=15) (**A, D**), Het-F (n=24) (**B, E**) and Het-U (n=16) (**C, F**), expressed as a ratio of their respective WT control values (denoted as a dotted line, n=21, 18 and 21, respectively), *P < 0.05 and ***P < 0.001 *versus* WT, unpaired t test. Data presented as means ± SEM.

### Fetal p110α is essential for embryonic development and trophoblast p110α is critical for its ability to up-regulate amino acid transport to match fetal demands for growth

We wanted to know more about the regulation of placental resource allocation to the developing fetus when there is a loss of fetal and trophoblast p110α. In particular, we wondered whether adaptation of placental transport function would still occur in heterozygous mutants (Het-U) if the trophoblast or fetal lineages were completely deficient in p110α. To do this, we selectively deleted the remaining p110α from the trophoblast or the fetal compartment of Het-U mice, using *Cyp19*Cre and *Meox2*Cre, respectively (termed Hom-P and Hom-F, respectively). We found that deleting the remaining p110α from the fetal lineages was lethal between days 11 and 12 of pregnancy (Table 1). In contrast to the lethality of Hom-F embryos, we found viable Hom-P fetuses in late gestation (Fig. 1C and Table S2). The timing of Hom-F lethality was identical to mutants with constitutive homozygous deficiency of p110α (α/α; (Foukas et al., 2006)) and is consistent with the role of p110α in early murine embryonic development (Xu et al., 2009). Taken together, these findings highlight that p110α in the fetal, but not the trophoblast compartment of the conceptus, is obligatory for prenatal development.

**Table 1.** Deleting the remaining p110α from fetal compartment of the conceptus results in fetal lethality in Hom-F mutants between days 11 and 12 of pregnancy. Frequency of viable fetuses in a litter per gestational age are displayed, with the number of litters in parentheses.

|  | WT | Het-F | Het-U | Hom-F |
| --- | --- | --- | --- | --- |
| D10 (5) | 23% | 26% | 34% | 17% |
| D11 (6) | 35% | 23% | 29% | 13% |
| D12 (2) | 50% | 25% | 25% | 0% |
| D13 (3) | 39% | 33% | 28% | 0% |

When comparing the growth of the Hom-P conceptuses to the control Het-U littermates, we found that fetal growth was restricted by a further 8% on day 19 of pregnancy (Table 2). However, despite the reduction in fetal growth, there was no difference in placental weight and labyrinthine morphology in Hom-P *versus* Het-U (Table 2). These observations suggest that the more severe reduction in fetal growth in Hom-P (relative to Het-U), was not caused by additional defects in the formation of the placental exchange region due to a complete loss of p110α in the trophoblast.

**Table 2.** Deleting the remaining p110α from the trophoblast reduces fetal weight but does not alter placental growth on day 19 of pregnancy, in Hom-P relative to Het-U controls. Conceptus weights are from n≥18, Lz and Jz volume from n≥6 and Lz morphology from n≥4 per genotype. Data are presented as means ± SEM. * *versus* Het-U, P<0.05, unpaired t test.

|  | Het-U | Hom-P |
| --- | --- | --- |
| Fetus (mg) | 1075±31 | 994±22* |
| Placenta (mg) | 84±3 | 89±3 |
| Labyrinthine zone (mm<sup>3</sup>) | 38±3 | 36±2 |
| Junctional zone (mm<sup>3</sup>) | 31±3 | 30±2 |
| Maternal blood spaces (mm<sup>3</sup>) | 13±3 | 11±1 |
| Trophoblast (mm<sup>3</sup>) | 20±1 | 22±2 |
| Fetal capillaries (mm<sup>3</sup>) | 3±1 | 4±1 |
| Barrier thickness (μm) | 4.0±0.2 | 3.9±0.1 |
| Surface area (mm<sup>2</sup>) | 11±1 | 10±1 |

We wondered whether the greater reduction in fetal growth may have been caused by a defect in placental transport function (a failure of the placenta to adapt its transfer capacity) in the Hom-P *versus* the Het-U mutants. We found that Hom-P placentas transferred 30% less amino acid (MeAIB) for the surface area available than Het-U littermates (Fig. 5A). Furthermore, Hom-P fetuses received less MeAIB solute for their size, as well as overall (Fig. 5B and Fig. S4). As the Het-U placenta up-regulated its transport capacity (Fig. 4C), this suggests that the greater reduction in fetal growth was due to an inability of the Hom-P placenta to adaptively increase amino acid transfer to the fetus. Placental glucose transfer capacity however, was not affected by a lack of placental p110α; transfer of MeG by the Hom-P placenta was equal to Het-U (Fig. 5A-B and Fig. S4). Data on feto-placental growth and placental transfer in Hom-P compared to wild-type and Het-P, which in contrast, retain p110α in fetus, are shown in Tables S3 and S4. Collectively, our data suggest that the demand signals of the compromised feto-placental unit for more amino acids operate via p110α in the trophoblast.

**Fig. 5.**
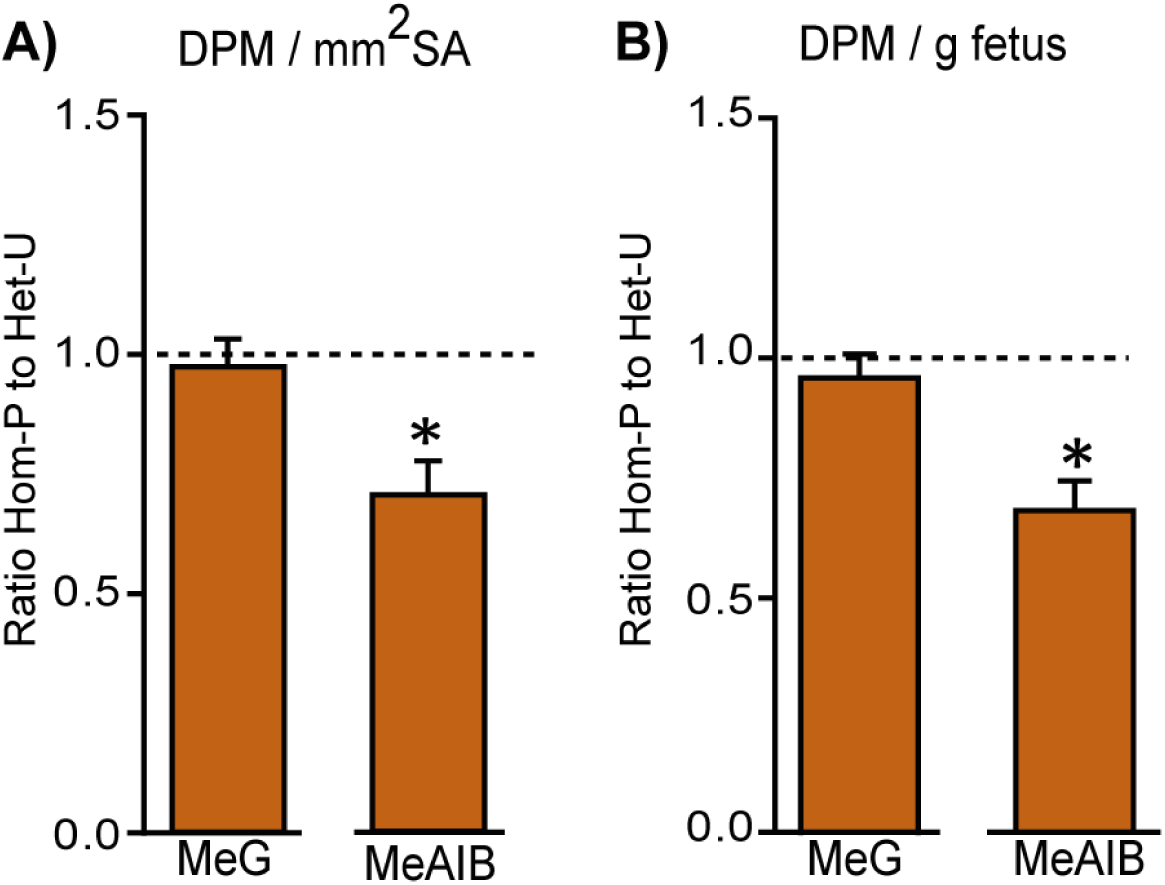
Retaining trophoblast p110α is critical for the ability of the placenta to increase amino acid transport. MeG and MeAIB transport relative to surface area available (**A**) or to fetal weight (**B**) on day 19 of pregnancy in Hom-P (n=16), expressed as a ratio of Het-U control values (denoted as a dotted line, n=16), *P < 0.05 *versus* Het-U, unpaired t test.

### Several genes downstream of p110α in the trophoblast are implicated in changes in placental phenotype to support the growing fetus

To identify genes responsible for the phenotypic differences observed between Het-U and Hom-P placentas at day 19 of pregnancy we compared their transcriptome using RNA-seq. We identified 97 differentially expressed genes, with 61 up- and 36 down-regulated in Hom-P *versus* Het-U placenta (Table S5). As expected, the FPKM (fragments per kilobase of transcript per million mapped reads) for the floxed exons, 18-19 of *Pik3ca* (Graupera et al., 2008) was diminished for Hom-P *versus* Het-U (Fig. S5). Using DAVID functional annotation followed by REViGO filtering we found that genes that were down-regulated between Het-U and Hom-P placentas have been implicated in cytolysis and cell death, proteolysis, regulation of immune effector processes, whilst those that were up-regulated have proposed roles in cellular hormone metabolic processes (Fig. 6A and Fig. S6). We confirmed using qRT-PCR the differential expression of several granzyme encoding genes, which are implicated in apoptosis (*Gzmc, Gzmf, Gzme, Gzmd, Gzmb* and *Gzmg*) (Table S5). However, paradoxically, we found increased levels of apoptosis in the junctional zone of the Hom-P relative to the Het-U placenta (Fig. 6B and C). Inspecting the list of genes from the RNA-seq dataset revealed that expression of the main glucose and System A amino acid transporters (*Slc2a, Slc38a*) and their proposed regulators (*Mtor, Tbc1d4*) (Miinea et al., 2005; Rosario et al., 2012) was not different between Hom-P and Het-U placentas. However, genes involved in water and oxygen transport (eg. *Aqp1, Aqp5, Hba-a2*) were altered (Table S5), suggesting that other transport capabilities of the placenta may be altered and contribute to the greater fetal growth restriction observed in the Hom-P *versus* Het-U. Other differentially expressed genes that were not featured in the pathway analysis have been implicated in regulating placental physiology (eg, *Cdx2, Cited2, Prl5a1, Prl7c1, Psg19* and *Psg22*) and pregnancy outcome (eg, *Fgl2, Acta2, Ngf* and *Nov/Ccn3*) (Table S5).

**Fig. 6.**
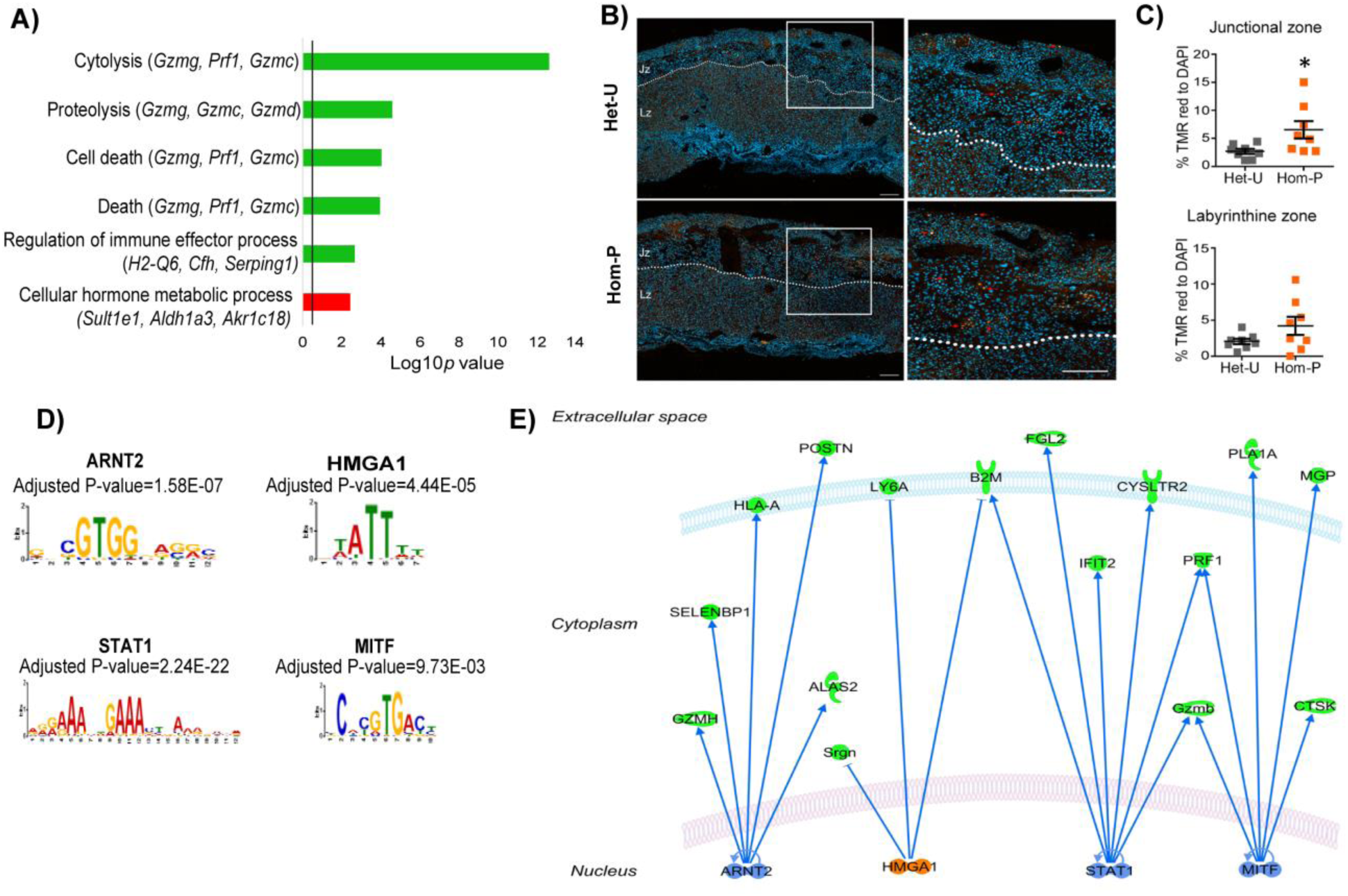
Genes downstream of p110α in the trophoblast implicated in changes in placental phenotype on day 19 of pregnancy. (**A**) Top-scoring biological processes enriched in Hom-P (n=4) *versus* Het-U (n=5) placentas on day 19 determined by RNA-seq (see also Supplementary Table S5). GO terms enriched in up-regulated genes shown in green and those down-regulated in red. Three genes with the highest fold expression changes are indicated in parentheses. The line corresponds to P=0.05. (**B**) Representative photomicrographs of increasing magnification of cells in the placenta undergoing apoptosis *in situ* in Het-U and Hom-P mutants on day 19 of pregnancy. Arrows indicate cells undergoing cell death (Tunnel: TMR red, nuclei: DAPI). (**C**) Quantification of cells in the placenta undergoing apoptosis *in situ* for Het-U (n=8) *versus* Hom-P (n=8) on day 19 of pregnancy. Scale bar=200μm, Tunnel: TMR red, nuclei: DAPI, Jz=Junctional zone, Lz=labyrinth zone. * P<0.05, unpaired t test. Data presented as means ± SEM. (**D**) Transcription factors with binding sites enriched at the promoters of differentially expressed genes, as identified by Analysis of Motif Enrichment (AME). (**E)** Regulatory network built with the four TFs identified by AME analysis using ingenuity pathway analysis. In green are shown proteins that are down-regulated at mRNA level in Hom-P versus Het-P placentae, in blue are depicted TFs predicted as repressors and in orange TFs predicted as activators.

We next identified significant enrichment of binding sites for four transcription factors (STAT1, ARNT2, HMGA1 and MITF) at the promoters of genes differentially expressed between Hom-P and Het-U placenta (Figure 6D and E). The mRNA expression of the transcription factors were not altered in the Hom-P versus Het-U placenta (Table S5), although previous work has shown that their activity is modified by PI3K signalling (Mounayar et al., 2015; Sheu et al., 2017; Terragni et al., 2011; Zhang et al., 2018) and/or implicated in the development of pregnancy complications related to placental dysfunction (Than et al., 2018). Therefore, p110α operates via several genes in the trophoblast to alter placental phenotype to support fetal growth.

### PI3K p110α loss differentially affects the expression of genes in the trophoblast and fetal cell lineages

We wanted to know whether the genes differentially expressed (DEGs) between Hom-P and Het-U placentas, were expressed in the trophoblast or embryonic lineage of the conceptus. Thus, we aligned our RNA-seq dataset with existing trophoblast and embryonic stem cell (TS and ES cell) transcriptomes using SeqMonk (Chrysanthou et al., 2018; Latos et al., 2015). We found that 46 of DEGs were expressed in both cell types, and of these, 26 were highly enriched in the TS compared to the ES cells (Fig. S7).

Functional analysis revealed that the DEGs enriched in TS cells have been implicated in the control of extracellular exosome, whereas those enriched in ES cells are involved in regulating gene expression and extracellular space (Table S6). These data suggested that p110α may modulate different types of genes in the trophoblast and fetal cell lineages of the conceptus and that some, but not all of the gene changes identified in our Hom-P *versus* Het-U analysis may be a direct consequence of p110α loss in the trophoblast. To explore these notions further, we deleted exons 18-19 of the *Pik3ca* gene using CRISPR/Cas9, to diminish p110α protein expression in TS cells (Fig. 7A-C). We then quantified the expression of a subset of DEGs identified *in vivo* in the mutant p110α and wild-type TS cells (Fig. 7D). These experiments were additionally performed in ES cells to assess if loss of p110α would have a similar effect on gene expression, as observed in mutant TS cells.

**Fig. 7.**
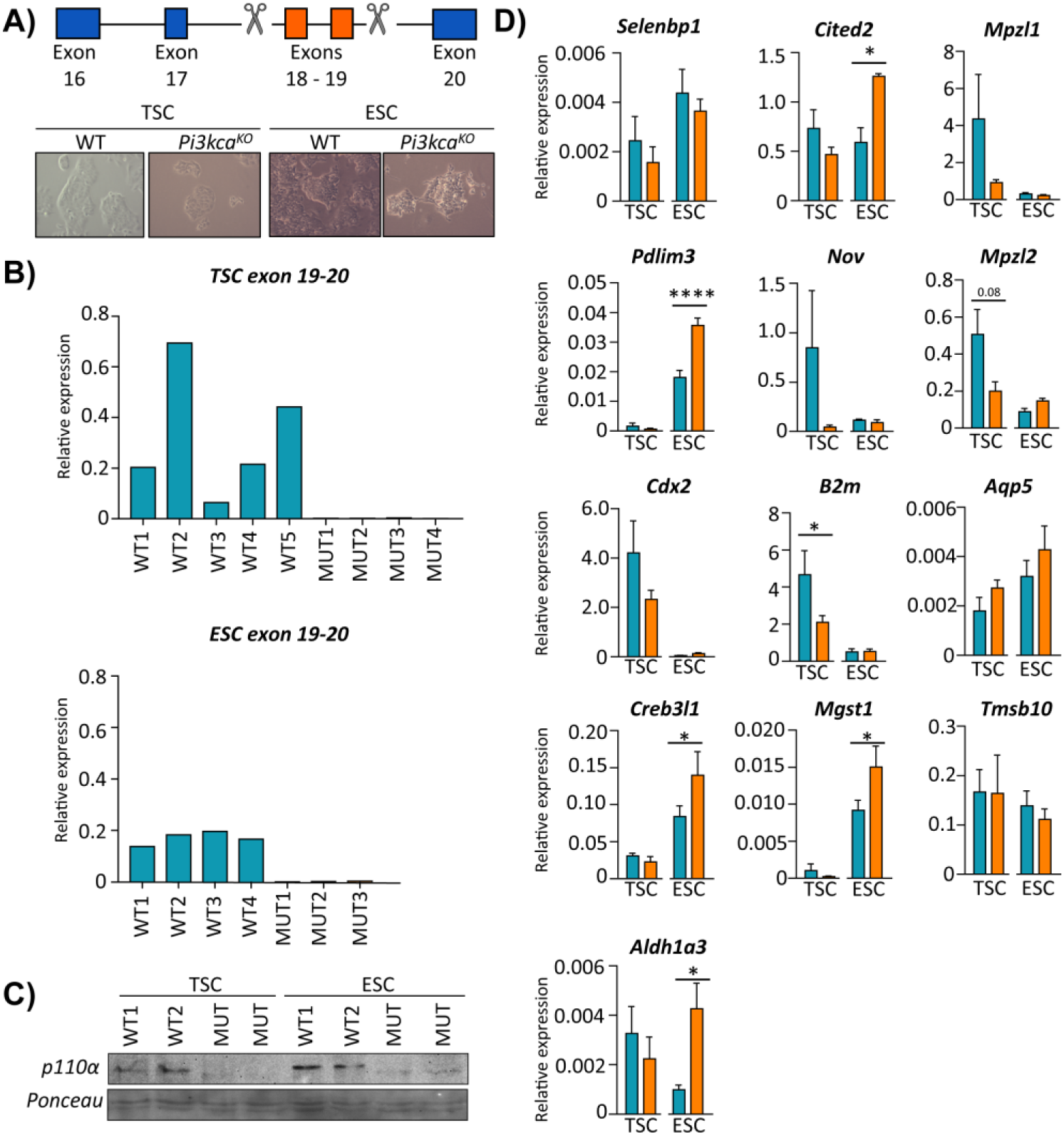
PI3K p110α differentially affects the expression of genes in the trophoblast and fetal cell lineages. (A) Details of CRISPR/Cas9 design to delete exons 18-19 of *Pik3ca* in trophoblast or embryonic stem cells (TSC and ESC, respectively). The deletion in TSC (trophoblast stem cells) and ESC (embryonic stem cells) was confirmed by (B) qRT-PCR analysis of exons 18-19 of *Pik3ca* and (C) by Western blotting for p110*α*. (D) The expression of candidate genes in *Pi3kca* wildtype (WT) and mutant (Mut) TSC and ESC. Data in (D) are normalized against housekeeping *Sdha*. * P<0.05, **** P<0.0001, two-ways ANOVA analysis followed by Fisher post hoc test. Data presented as means ± SEM.

Loss of p110α significantly reduced the expression of *B2m* in mutant TS cells (Fig. 7D), similar to findings *in vivo* from the Hom-P *versus* Het-U placenta (Table S5). We also observed a trend for reduced expression of *Mzp12* upon TS p110α loss, although this did not reach statistical significance with the small number of biological replicates used (P=0.079; Fig. 7D). However, none of the other DEGs analyzed were affected by TS p110α deficiency. These data suggest that the mis-regulated pattern of genes identified in the Hom-P *versus* Het-U placenta represent a combination of both direct and secondary effects of trophoblast p110α loss.

Loss of p110α also had a more prominent impact on the expression of DEGs in ES cells. For instance, p110α deficiency significantly increased the expression of *Cited2, Pdlim3, Creb3l1, Mgst1, Aldh1a3* in the ES cells (Fig. 7D). For wildtype clones, the expression of the *Pik3ca* was overall, more uniform in ES than TS cells (Fig. 7B). Moreover, loss of p110α tended to increase the expression of genes analyzed in ES cells, but decreased them in TS cells. Taken together, these data demonstrate that pathways downstream of p110α differentially affect the expression of genes in the trophoblast and fetal cell lineages of the conceptus.

### Discussion

Here, using *in vivo* conditional genetic manipulations of p110α we have identified how the trophoblast and fetal cell lineages interact to control placental resource supply and thereby, affect healthy growth of the fetus. The major cause of intrauterine growth restriction is placental insufficiency in the developed world (Baschat and Hecher, 2004). Our results are therefore, important for understanding the etiology of placental insufficiency and intrauterine growth restriction.

The data presented here indicate that p110α in the fetal, but not the trophoblast compartment of the conceptus determines placental size in late mouse pregnancy. The mechanism by which loss of fetal p110α reduces formation of the placenta is unknown. Elongation of the fetal capillary network is required for growth of the placenta, particularly its labyrinthine zone (Adamson et al., 2002; Coan et al., 2004). Reduced fetal capillary formation in the labyrinthine zone of the Het-F could therefore underlie the reduced placental weight and labyrinthine zone observed and data are consistent with previous work showing defective angiogenesis and vascular development in the fetus and placenta in response to p110α loss (Graupera et al., 2008; Sferruzzi-Perri et al., 2016). The reduction in fetal and placental weight was similar for Het-F conceptuses (both reduced by ∼11%), suggesting that perhaps the placenta grew to match the genetically-determined growth requirements of the fetus (Sandovici et al., 2012). However, it is also plausible that placental growth decreased to match the experimentally induced slower growth of the Het-F fetuses. Herein, the direct effects of fetal p110α cannot be isolated from the indirect effects on the fetal growth restriction in Het-F on day 19 of pregnancy. Thus, future work should assess the ontogeny of changes in placental phenotype with respect to the growth of the fetus in the Het-F during gestation.

The lack of an effect of trophoblast p110α deficiency in Het-P and Hom-P on placental weight was surprising as both loss of IGF2 and protein kinase B (AKT), which are main up- and down-stream mediators of PI3K signaling respectively cause placental growth restriction (Constancia et al., 2002; Kent et al., 2012; Plaks et al., 2011; Yang et al., 2003). Moreover, studies using non-isoform specific inhibitors have shown that Class 1A PI3Ks mediate the proliferative and survival effects of ligands on trophoblast *in vitro* (Diaz et al., 2007; Forbes et al., 2008). Deleting p110α using the conditional *Pik3ca* line in the current study does not alter the stoichiometry of PI3K complexes (Graupera et al., 2008), and thus it is unlikely that other PI3K isoforms compensate for the loss of p110α in the trophoblast. In the current study however, the expression of cell death genes were down-regulated and the level of apoptosis in the Hom-P compared to Het-U placenta paradoxically upregulated on day 19 of pregnancy (∼1 day prior to term). Differences between gene expression and tissue levels of apoptosis in our study may be relate to the low volume density (∼20%) of the junctional zone in the whole placenta which was used in the RNAseq analysis. Assessing the ontogeny of changes in cell death genes and the activation of apoptosis in the Hom-P placenta is required. Moreover, whether placental weight (as well as morphology) may be decreased in Hom-P *versus* Het-U in the last 24h prior to delivery remains to be determined.

Our stereological analyses of the Het-F, Het-P and Het-U placentas demonstrated that epiblast-derived and trophoblast p110α have non-overlapping contributions in regulating the development of the placental exchange region. In particular, p110α in the epiblast-derived compartment determined the size of the placental labyrinthine zone and its trophoblast constituent, whereas p110α in the trophoblast was important for expanding the surface area for maternal-fetal exchange. Moreover, p110α in both the trophoblast and epiblast-derived lineages contributed to the formation of the capillary network and the thinness of the diffusion barrier in the placental exchange region. These findings are consistent with previous work demonstrating that the PI3K pathway regulates trophoblast differentiation (Kent et al., 2010) and defects in p110α signaling impair developmental angiogenesis and placental exchange region morphogenesis (Graupera et al., 2008; Lelievre et al., 2005; Sferruzzi-Perri et al., 2016). As fetal capillary volume was affected by a deficiency of p110α signaling in the trophoblast, and trophoblast volume was altered by a deficiency in fetal p110α, our data suggest that there is a paracrine communication between cellular compartments in the placental labyrinthine zone, which occurs via p110α signaling. Paracrine signaling via p110α may also explain why when comparing the three heterozygote lines, the Het-P placenta uniquely showed less maternal blood spaces and a greater trophoblast volume. Paracrine communication between trophoblast and endothelial cell lineages in the labyrinthine zone has been reported previously for other signaling proteins (Limbourg et al., 2005; Lu et al., 2013; Moreau et al., 2014; Tang et al., 2006). Future work should evaluate the density of different trophoblast cell types in the labyrinthine zone (eg. syncytial trophoblast layer 1, syncytial trophoblast layer 2 and sinusoidal giant cells) in the Het-P and other p110α mutant placentas.

Our analysis of placental transfer function *in vivo* revealed that independent of specific changes in placental labyrinthine morphology with fetal *versus* trophoblast p110α loss, fetal glucose and System A-mediated amino acid supply relative to the size of the fetus, was maintained or even increased due to an adapting placenta which increased materno-fetal substrate clearance per unit surface area. We found that in conceptuses with p110α deficiency, this adaptive up-regulation of placental amino acid transport depended on the expression of p110α in the trophoblast; System A-mediated amino acid transport was significantly lower for the Hom-P compared to the adapted Het-U placenta. This inability of the Hom-P placenta to up-regulate System A amino acid transport, may explain, at least partly, the further reduction in fetal weight, when compared to Het-U. Indeed, inhibiting System A-mediated amino acid transport during rat pregnancy *in vivo* leads to fetal growth restriction (Cramer et al., 2002) and reductions in System A transporter activity occur prior to the onset of intrauterine growth restriction in rats and baboons (Jansson et al., 2006; Pantham et al., 2016). Previous work has shown that constitutive heterozygous loss of p110α activity in the conceptus (α/+ mutants) does not affect the expression of glucose (*Slc2a*) or the sodium coupled System A amino acid transporter (*Slc38a*) genes in the placenta (Sferruzzi-Perri et al., 2016). We also did not find differential expression of these transporters in Hom-P compared to Het-U placenta by RNA-seq. In other cell types, p110α alters the activity of glucose and sodium transporters on the plasma membrane (Frevert and Kahn, 1997; Katagiri et al., 1996; Wang et al., 2008). Loss of p110α may therefore, alter glucose and amino acid transport to the fetus by the placenta *via* post-transcriptional mechanisms. Previous work has shown that when placental supply and fetal demand for resources to grow are mis-matched, IGF2 in the fetus may directly or indirectly signal demand to the placenta to adaptively up-regulate its transport of glucose and amino acids (12, 14, 26, 57). Our data therefore imply that demand signals of the compromised feto-placental unit, such as IGF2, operate via p110α in the trophoblast to adapt amino acid supply to the fetus for growth. However, it is equally possible that fetal demand signals operate via pathway/s independent of p110α. Moreover, it is plausible that these pathway/s may be sufficient to compensate for the effects of moderately down-regulated p110α in the Het-P, but are insufficient when p110α is markedly knocked-down as in the Hom-P. Further work is required to identify the elusive fetal demand/s signals and explore the contribution of other signaling pathways implicated in environmental sensing, in driving alterations in placental transport phenotype in response to p110α deficiency, including the mechanistic target of rapamycin (mTOR), mitogen-activated pathway (MAPK), general control nonrepressed 2 (GCN2), glucokinase, G-protein coupled-receptors (GPCRs) and adenosine monophosphate-activated protein kinase (AMPK) (Efeyan et al., 2015).

Our transcriptomic comparison of the Hom-P and Het-U placentas identified several genes downstream of p110α that may be important in adapting placental phenotype to support fetal growth. Of note, we found 6 granzyme encoding genes, to be down-regulated in expression between Hom-P and Het-P. Others have shown that the PI3K pathway regulates granzyme expression and function of different leukocyte populations (Blanco et al., 2015; Efimova and Kelley, 2009; Jiang et al., 2000), and thus p110α could be playing a similar role in trophoblast. We also found that some of the genes differentially expressed between Hom-P and Het-U placenta have been previously implicated in the regulation of trophoblast differentiation (eg, *Cdx2*) (Strumpf et al., 2005), growth and transport (*Cited2*)(Withington et al., 2006) and hormonal regulation of maternal physiology (*Prl5a1, Prl7c1, Psg19* and *Psg22*) (Moore and Dveksler, 2014; Soares et al., 2007). Moreover, a number of genes identified have been implicated in miscarriage (*Fgl2*) (Clark et al., 2001) and in the development of placental dysfunction in the pregnancy complication, preeclampsia (*Acta2, Ngf* and *Nov/Ccn3*)(Gellhaus et al., 2006; Sahay et al., 2015; Todros et al., 2007). We found significant enrichment of binding motifs for the transcription factors, STAT1, ARNT2, HMGA1 and MITF in the promoters of genes differentially expressed between Hom-P and Het-U. Others have demonstrated that the PI3K pathway modulates the activity of three of these transcription factors (Mounayar et al., 2015; Sheu et al., 2017; Terragni et al., 2011; Zhang et al., 2018) and of note, there is an interaction between p110α and STAT1 activity (Mounayar et al., 2015). Many of the differentially expressed genes in Hom-P *versus* Het-U placenta, were found to be enriched in TS cells however of the fourteen candidates studied, only one was significantly altered in response to TS cell p110α deficiency. Thus, the mis-regulated pattern of genes identified in the Hom-P *versus* Het-U placenta may be both a direct consequence of p110α in the trophoblast, as well as the result of indirect effects on the embryonic lineages (such as fetal capillaries) in the placenta, or in the decidua. Single-cell RNA-seq should be performed on p110α mutant placentas in the future to substantiate this interpretation. Further work is additionally required to assess more directly the contribution of genes differentially expressed between Hom-P and Het-U placentas, in adaptive responses of the placenta (Sferruzzi-Perri et al., 2011).

Loss of p110α in ES cells also affected the expression the candidate genes studied. However, there were many more candidate genes significantly altered in the mutant ES cells and the pattern of change was largely distinct to that observed the mutant TS cells. Indeed, genes were typically upregulated in ES cells, but reduced in TS cells in response to p110α deficiency. Moreover, the genes differentially expressed between Hom-P and Het-U and enriched in TS and ES cells were predicted to participate in diverse cell functions, for example extracellular exosome and gene expression, respectively. These data imply that p110α regulates the expression of different genes and functional pathways in the trophoblast and epiblast-derived tissues of the conceptus. They also reinforce that the embryonic lineages of the conceptus are more sensitive to p110α loss than the trophoblast, and may provide some explanation for the lethality observed for the Hom-F but not Hom-P mutants. These findings could be furthered in the future by performing RNAseq on TS and ES cells to obtain a complete transcriptomic comparison in response to p110α deficiency.

Previous work has shown that the genotype and environment of the mother modulates placental growth and transport function (Angiolini et al., 2011; Constancia et al., 2005; Sferruzzi-Perri et al., 2016; Sferruzzi-Perri et al., 2017; Sferruzzi-Perri et al., 2011). In the current study, all data were obtained from litters of mixed genotype and only comparisons to their respective control siblings were made to minimize possible effects of the maternal metabolic environment on placental phenotype. However, changes in placental phenotype with fetal and trophoblast *Pik3ca* deletion can still occur within the homogenous maternal metabolic environment and according to fetal genotype, which was not assessed in the current study. Future work should therefore, attempt to explore the interaction between maternal p110α metabolic profile and fetal genotype in the regulation of placental resource allocation. This can be done initially by measuring maternal hormones and glucose/amino-acid levels in the diverse fetal/maternal genotype combinations, but quantification of the ‘individual’ effects will be highly challenging.

Collectively, our data indicate that p110α plays distinct roles in the different compartments of the conceptus, which ultimately affect placental phenotype and fetal growth. In particular, p110α in the fetal lineages of the conceptus is essential for prenatal development and a major regulator of placental phenotype (Fig. 8). Moreover, trophoblast p110α signaling is critical for its ability to up-regulate amino acid transport to match fetal demands for growth near term. However, the Hom-F and Hom-P have a heterozygous deficiency of p110α in the trophoblast and epiblast-derived lineages, respectively. Thus, there may be interactions between the different compartments of the conceptus that are heterozygous and homozygous deficient in p110α, which may have contributed to the placental and fetal phenotypes observed for Hom-F and Hom-P. Furthermore, gene disruption may occur up to 1.5 days earlier using the *Meox2*Cre, compared to the *Cyp19*Cre (at around days 5 and 6.5, respectively) (Tallquist and Soriano, 2000; Wenzel and Leone, 2007). Moreover, the *Cyp19*Cre exhibits mosaic activity (observed herein and reported previously (Wenzel and Leone, 2007)). Therefore, temporal differences in the timing of Cre activation and levels of Cre activity between the *Meox2*Cre and *Cyp19*Cre lines may have contributed to feto-placental phenotypes observed for the fetal *versus* trophoblast p110α mutants, in the current study. In addition, previous characterization of *Meox2*Cre line has shown Cre is active in the yolk sac mesoderm (Tallquist and Soriano, 2000). As the yolk sac is essential for fetal growth, loss of p110α from the yolk sac mesoderm could have contributed to the fetal phenotypes observed with the Het-F and Hom-F. Employing alternative strategies to target expression of p110α in different conceptus lineages, such as using lentiviral-mediated shRNA/siRNA transfection of mouse blastocysts (Georgiades et al., 2007; Okada et al., 2007) are therefore, warranted in the future. Nevertheless our findings are important in the context of human pregnancy, as dysregulated expression of both up and down-stream components of the PI3K pathway in the placenta are associated with abnormal fetal growth and poor pregnancy outcome (reviewed in (Sferruzzi-Perri et al., 2017)).

**Fig. 8.**
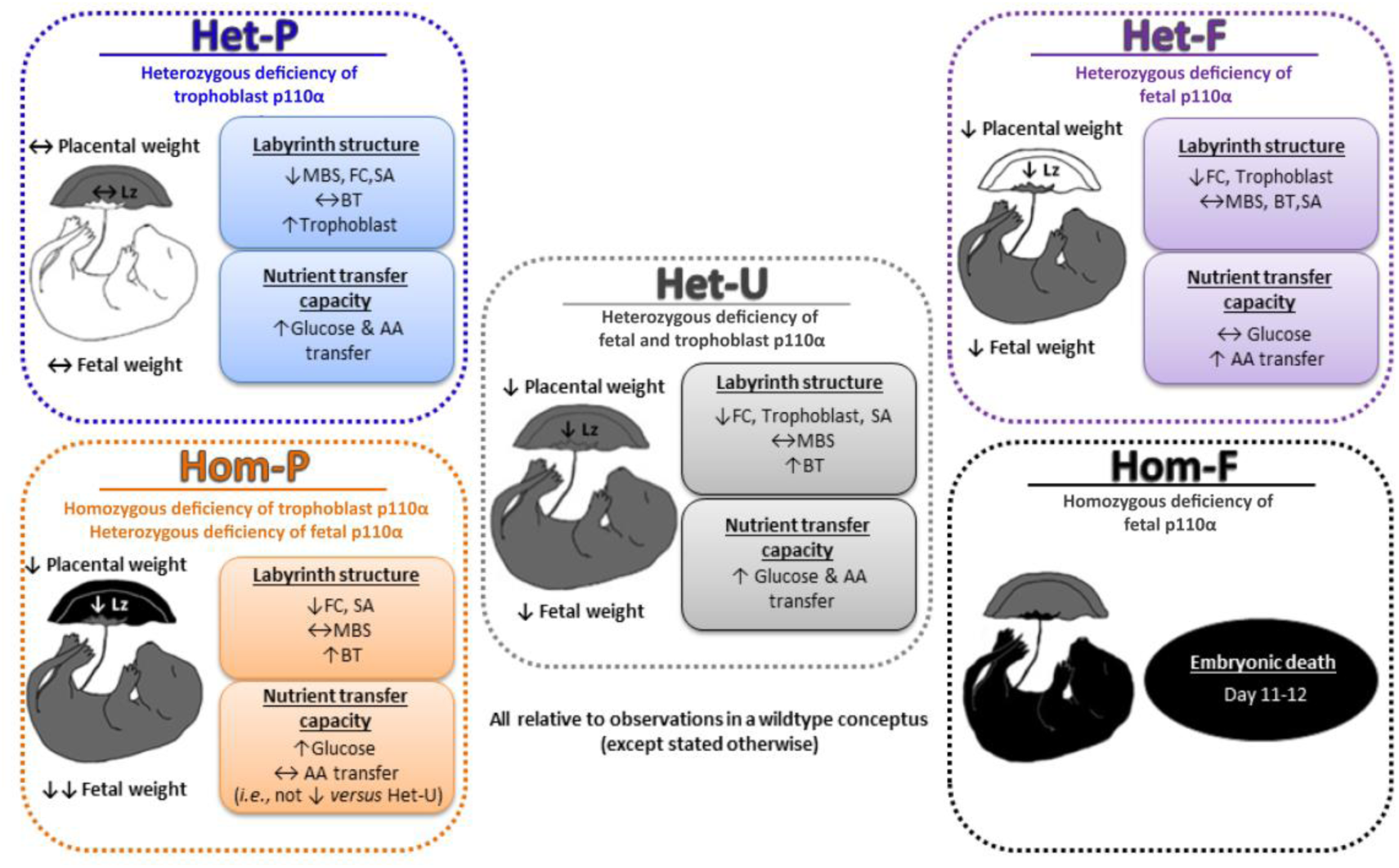
Summary figure showing the phenotypes of all p110α mutants used in this study, which together highlight that fetal and trophoblast p110α have distinct contributions in regulating resource allocation to the growing fetus. Findings are from mice at day 19 of pregnancy, unless indicated otherwise. AA=amino acid, BT=barrier thickness, FC=fetal capillaries, Lz=labyrinth zone, MBS=maternal blood spaces, SA=surface area for exchange.

## Materials and Methods

### Mice and genotyping

Mice were housed under dark:light 12:12 conditions with free access to water and chow in the University of Cambridge Animal Facility abiding by the UK Home Office Animals (Scientific Procedures) Act 1986 and local ethics committee. Mice in which exons 18 and 19 of the p110α gene were flanked by LoxP sites (*Pik3ca*^*Fl*^) on a C57Bl6 background were kindly provided by Dr Bart Vanhaesebroeck (University College London) and Dr Klaus Okkenhaug (Babraham Institute) (Graupera et al., 2008). These were time-mated with those expressing the Cre recombinase gene under the control of the trophoblast-specific human *Cyp19* promoter (Wenzel and Leone, 2007), the embryonic *Meox2* promoter (Tallquist and Soriano, 2000) or the ubiquitous *CMV* promoter (Schwenk et al., 1995). This achieved three models that had a ∼50% decrease in p110α in the trophoblast, fetal lineages or whole conceptus, termed Het-P, Het-F and Het-U, respectively. Further crosses were performed on a second generation of Het-U mice to selectively delete the remaining p110α allele in the trophoblast using *Cyp19*Cre (Hom-P) or fetal lineage using *Meox2*Cre (Hom-F). See Table S1 for experimental crosses used. Note the wild-types for the Het-P and Het-U are the same animals and the Het-U were the controls for Hom-P. The genotypes of the fetuses/mice were determined by PCR using primers to detect *Cyp19*Cre (Fwd: GACCTTGCTGAGATTAGATC and Rev: AGAGAGAAGCATGTTTAGCTGG), *Meox2*Cre (Fwd: GGACCACCTTCTTTTGGCTTC, Rev: AAGATGTGGAGAGTACGGGGTAG, Cre: CAGATCCTCCTCAGAAATCAGC) and *CMV*Cre (Cre Fwd: CGAGTGATGAGGTTCGCAAG, Cre Rev: TGAGTGAACGAACCTGGTCG, Internal control Fwd: ATGTCTCCAATCCTTGAACACTG and Internal control Rev: GCAGTGGGAGAAATCAGAACC). This was done using the following cycle conditions: 94°C, 3 min, then 35 cycles of 94°C for 30s, 57°C for 30s and 72°C for 90s, and then 72°C for 7min. Genotyping was also performed by PCR using primers for *Pik3ca*^*Fl*^ (Fwd:CTAAGCCCTTAAAGCCTTAC, Rev:CAGCTCCCATCTCAGTTCA and Deletion: ACACACTGCATCAATGGC) as described previously (Graupera et al., 2008) and by real time PCR (qRT-PCR) using Taqman probes (*Pik3ca:* Mm00435672_g1 and *Gapdh*: Mm99999915_g1) to confirm deletion in whole fetal and placental tissue homogenates. Due to the mosaic activity of the *Cyp19*Cre (Wenzel and Leone, 2007) a cut-off for *Pik3ca* deletion in the placenta using qRT-PCR was set for <65% Het-P and <30% for Hom-P.

To visualize the Cre recombinase activity, *Cyp19*Cre and *Meox2*Cre mice were mated to the *Rosa26*TdRFP reporter mouse line (*Gt(ROSA)26Sor;* purchased from the Jackson lab and a generous gift from Prof William Colledge, University of Cambridge) (Soriano, 1999). Fetuses and placentas were collected on day 16 of pregnancy, fixed in 4% (wt/vol) PFA, 30% (wt/vol) sucrose and then frozen in OCT (optimal cutting temperature media, TissueTek) for cryosectioning and staining in DAPI.

### Placental nutrient transfer assays

The unidirectional materno-fetal clearance of the non-metabolisable radioactive tracers, ^14^C-methyl amino-isobutyric acid (MeAIB) and ^3^H-methyl D-Glucose (MeG) were measured under anaesthesia with fentanyl-fluanisone (hypnorm):midazolam (hypnovel) in sterile water (1:1:2, Jansen Animal Health) (Sibley et al., 2004). A 200μl bolus containing 3.5 µCi, of MeAIB (NEN NEC-671; specific activity 1.86GBq/mmol, Perkin Elmer, USA) and 3.5 µCi MeG (NEN NEC-377; specific activity 2.1GBq/mmol) in physiological saline (0.9% wt/vol) was injected into the maternal jugular vein. Two minutes after tracer injection, the dam was killed by cervical dislocation, uteri were collected and fetal and placental weights recorded. Entire litters of placentas were collected for morphological analyses or snap frozen in liquid nitrogen for quantification of gene expression. Fetuses were decapitated, entire fetal tails taken for DNA genotyping and then fetuses minced and lysed at 55°C in Biosol (National Diagnostics, Atlanta, USA). Fetal lysates were measured for beta emissions by liquid scintillation counting (Optiphase Hisafe II and Packard Tri-Carb, 1900; Perkin-Elmer USA) and radioactivity (DPM) in the fetuses was used to calculate transfer relative to the estimated placental surface area or to fetal weight.

### Placental morphological analyses

Placentas were bisected and one half was fixed in 4% (wt/vol) paraformaldehyde, paraffin-embedding, exhaustively sectioned at 7 μm and stained with hematoxylin and eosin to analyse gross placental structure. The other half was fixed in 4% (wt/vol) glutaraldehyde, embedded in Spurr’s epoxy resin and a single, and 1-μm midline section was cut and then stained with toluidine blue for detailed analysis of labyrinthine zone (Lz) structure using the Computer Assisted Stereological Toolbox (CAST v2.0) program as previously described (Coan et al., 2004). Briefly, to determine the volume densities of each Lz component (fetal capillaries [FC], maternal blood spaces [MBS], trophoblast), point counting was used and their densities were multiplied by the estimated volume of the Lz to obtain estimated component volumes. Surface densities of maternal-facing and fetal-facing interhaemal membrane surfaces were then determined by counting the number of intersection points along cycloid arcs in random fields of view. These were converted to absolute surface areas and the total surface area for exchange calculated (averaged surface area of MBS and FC). The mean interhaemal membrane thickness was determined by measuring the shortest distance between FC and the closest MBS at random starting locations within the Lz. Per Lz, 200 measurements were made. The harmonic mean thickness (barrier thickness) was calculated from the reciprocal of the mean of the reciprocal distances.

### Placental in situ cell death staining

Cells undergoing cell death were detected in paraffin-embedded sections of placenta using the *In Situ* Cell Death Detection Kit, TMR red (Sigma Aldrich), according to the manufacturer’s instructions. The proportion of TMR red to DAPI stained cells was determined in each placental section using Image J (freeware software).

### Placental immuno-localisation of p110 α expression

Placental sections were washed with PBS to remove OCT and underwent antigen retrieval with citrate buffer before immunolabelling for p110α (1:75 dilution, Cell Signaling, C73F8). Sections were treated with 0.5% Triton X-100 before immunolabelling. Bound antibody was detected using biotinylated goat anti-rabbit IgG (Abcam, ab6720) followed by streptavidin-conjugated horseradish peroxidase (Rockland, S000-03) and Sigma fast DAB tablets according to manufacturer instructions. Sections were lightly counterstained with hematoxylin and mounted in DPX. Negative controls were performed by omission of the primary antibody.

### Placental RNA sequencing (RNA-seq)

Total RNA was extracted from whole placental halves using the RNeasy Plus Mini Kit (Qiagen, UK) according to manufacturer’s instructions. RNA-seq libraries were prepared using TruSeq (Illumina) and the barcoded libraries were combined and submitted for sequencing (50bp single ended on a HiSeq2500) by the Wellcome Trust-MRC Institute of Metabolic Science Genomics Core Facility. The number of reads per library was between 14.3 and 33.4 million. Fastq files were aligned to mouse genome GRCm38 with TopHat (version 2.0.11; Kim 2013) and default settings (The mapping percentage varied between 96.2 and 98.3%). Differential expression was performed using Cuffdiff (version 2.1.1; Trapnell 2012) with the following settings; require a minimum of 5 alignments to conduct significance testing, perform bias correction, do initial estimation to weight multi-hits. Genes identified as differentially expressed (DEG) with Cuffdiff significance indicated as q value<0.05 and 1.5-fold expression change. Pathway analysis was then determined using DAVID gene ontology (Huang da et al., 2009) and redundant GO terms were filtered-out using REViGO (Supek et al., 2011).

To search for enrichment of transcription factor binding sites at the promoters of DEG, we used Eukaryotic Promoter Database (EPD – https://epd.vital-it.ch/index.php) to retrieve the DNA sequences from 1,000bp upstream to 100bp downstream of the transcriptional start site. These sequences were then analysed using Analysis of Motif Enrichment (AME v4.12.0 – http://meme-suite.org/tools/ame) by selecting *Mus musculus* and HOCOMOCO Mouse (v11 FULL) as motif database. Transcription factors predicted by AME and Ingenuity Pathway Analysis (IPA) were then used for network visualization, performed using IPA.

### CRISPR/Cas9 knockout in Trophoblast and Embryonic Stem Cells

For CRISPR/Cas9-mediated deletion of *Pik3ca, E14tg2a* embryonic stem cells and *TS-Rs26* trophoblast stem cells (a gift of the Rossant lab, Toronto, Canada) were transfected with Cas9.2A-eGFP plasmid (Plasmid 48138 Addgene) harbouring guide RNAs flanking the exon 18 and exon 19 (http://CRISPR.mit.edu) (Fig. 7). Transfection was carried out with Lipofectamine 2000 (ThermoFisher Scientific 11668019) reagent according to the manufacturer’s protocol. Knockout clones were confirmed by western blotting and by genotyping using primers spanning the deleted exons, and by reverse transcription followed by semi-quantitative PCR (RT-qPCR) with primers (AGGGAGCACAAGAGTACACCA and GGCATGCTGCCGAATTGCTA) within and downstream of the deleted exons, as shown in Fig. 7. Values were normalized to *Sdha* expression. Three to five independent knockout clones were analysed for each cell line. The expression of fourteen genes found to be differentially regulated between Het-U and Hom-P placenta, were then assessed by qPCR in p110α deficient TS and ES cells.

### Gene expression by qRT-PCR

Total RNA was extracted from whole placental halves using the RNeasy Plus Mini Kit (Qiagen, UK) whereas total RNA was extracted from TS and ES cells using trizol (Invitrogen), chloroform and ethanol extraction. Multiscribe Reverse Transcriptase and random primers (Applied Biosystems) were used to synthesize cDNA from 2.5ug of RNA. Samples were analysed in duplicate by qRT-PCR (7500 Fast Real-Time PCR System, Applied Biosystems, UK) using Sybr green chemistry and pairs of forward and reverse primers (Table S7). The expression of genes of interest were normalised to the expression of *Gapdh* or *Sdha* which was not affected by genotype. Data were analysed using the 2–ΔΔCT method for quantification.

### Western Blotting

Protein expression was quantified using p110α antibody (1:500; Cell Signalling, 4249) as described previously (Sferruzzi-Perri et al., 2011).

### Statistics

Data are presented as mean ± SEM. Data were analysed using unpaired and paired t tests with Excel, as required or two-way ANOVA with GraphPad Prism 7. Data were considered statistically significant when P<0.05.

## Supporting information

## Acknowledgments

We thank the Centre for Trophoblast Research for the award of a Next Generation Fellowship to A.N.S.-P., and the Erasmus Exchange scheme and COST ACTION (EPICONCEPT and SALAAM) grants to J. L.-T. We thank Professor Abigail Fowden for allowing us to perform the animal procedures under her project licence and the staff at the Combined Animal Facility for their technical help. We thank Dr Bart Vanhaesebroeck and Dr Klaus Okkenhaug for providing the p110α-floxed mice and Professor William Colledge for the *tdTomato* mice. We thank Brian Y.H. Lam from Genomics/Transcriptomics Core Facilities, MRC Metabolic Diseases Unit, for help with preliminary RNA-Seq data analysis and data submission to public repositories. B.Y.H.L. is supported by MRC grant (MRC_MC_UU_12012/5). Finally, we thank Dr Jacqui Sheids and Dr Angela Riedel for scanning our fluorescently labelled slides and technical support.

## Author contributions

A.N.S.-P. and M.C. designed the research. J.L-T.,V.P-G., J.K., L.C.K, W.N.C., A.A., I.G., E.F.L., M.H., I.S., and A.N.S-P., performed the research and analyzed the data. A.N.S.-P. and J.L-T., wrote the manuscript which was revised and approved by all other authors. A.N.S-P. and M.C. supervised the research.

## Data and materials availability

The RNA-seq data is currently being deposited at the University of Cambridge data repository and the accession number will soon be provided.

## Supplementary Materials

**Fig. S1.**
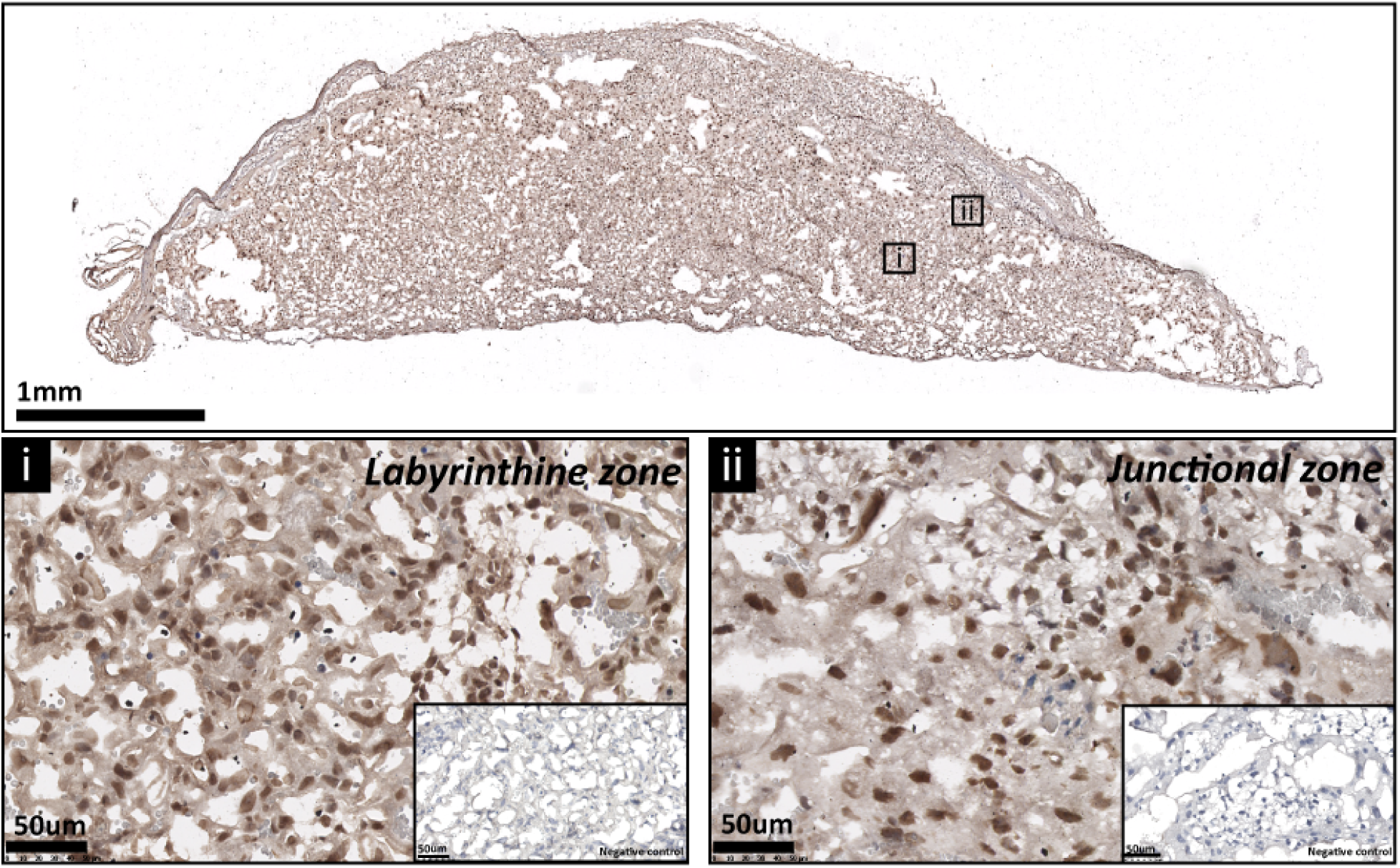
The expression of p110α protein by the mouse placenta on day 19 of pregnancy. Representative stained section shown with negative control also displayed.

**Fig. S2.**
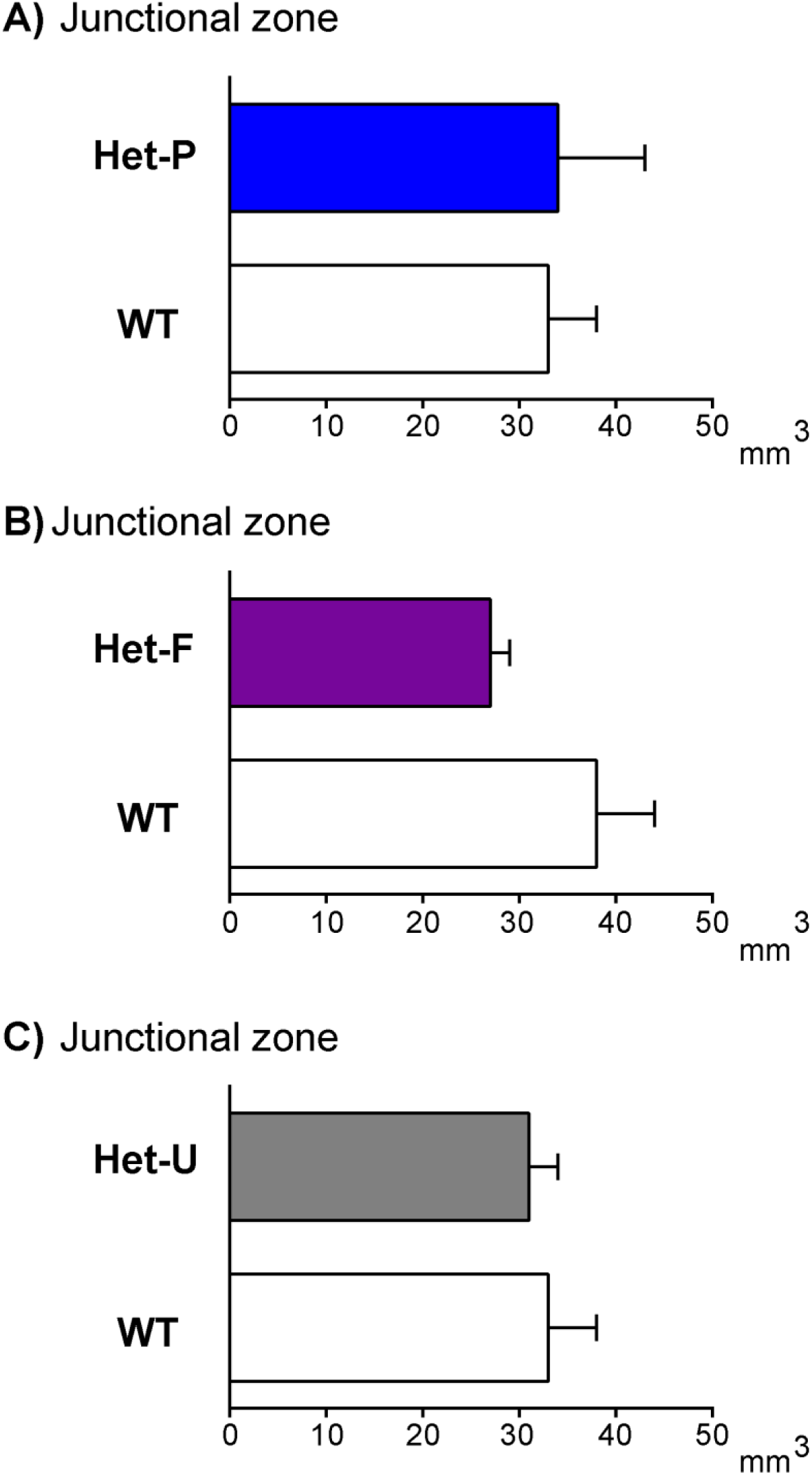
Junctional zone volume is not modified by fetal and/or trophoblast loss of p110α. Placental junctional zone volume on day 19 of pregnancy in Het-P (**A**), Het-F (**B**) and Het-U (**C**). n≥6 placentas were assessed for each genotype. Data are presented as means ± SEM.

**Fig. S3.**
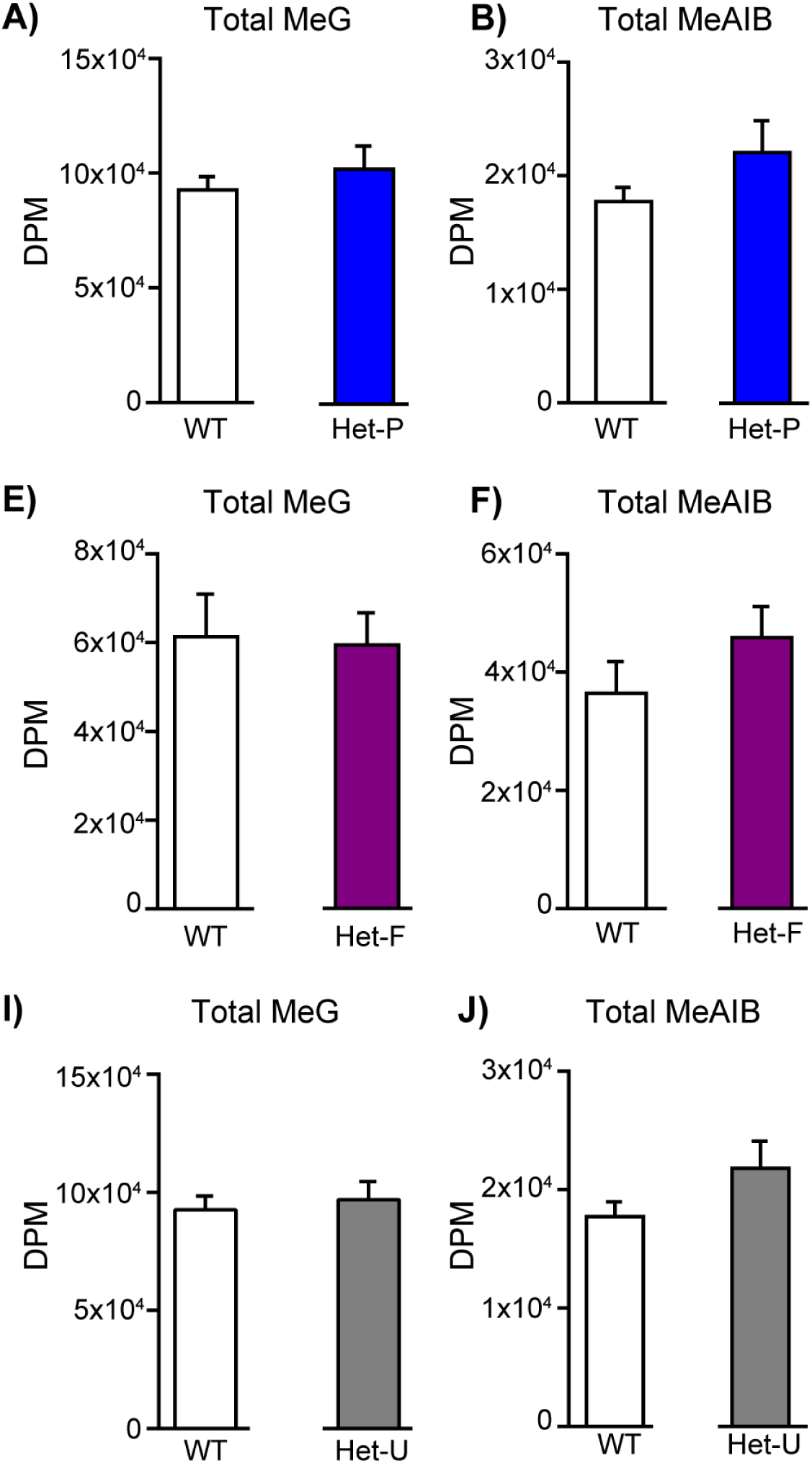
Total solute accumulation in response to fetal and/or trophoblast loss of p110α. The accumulation of ^3^H-methyl-D glucose (MeG) and ^14^C-amino isobutyric acid (MeAIB) on day 19 of pregnancy for Het-P (n=15) (**A, D**), Het-F (n=24) (**B, E**) and Het-U (n=16) (**C, F**) with their respective WT control values (n=21, 18 and 21, respectively), Data presented as means ± SEM.

**Fig. S4.**
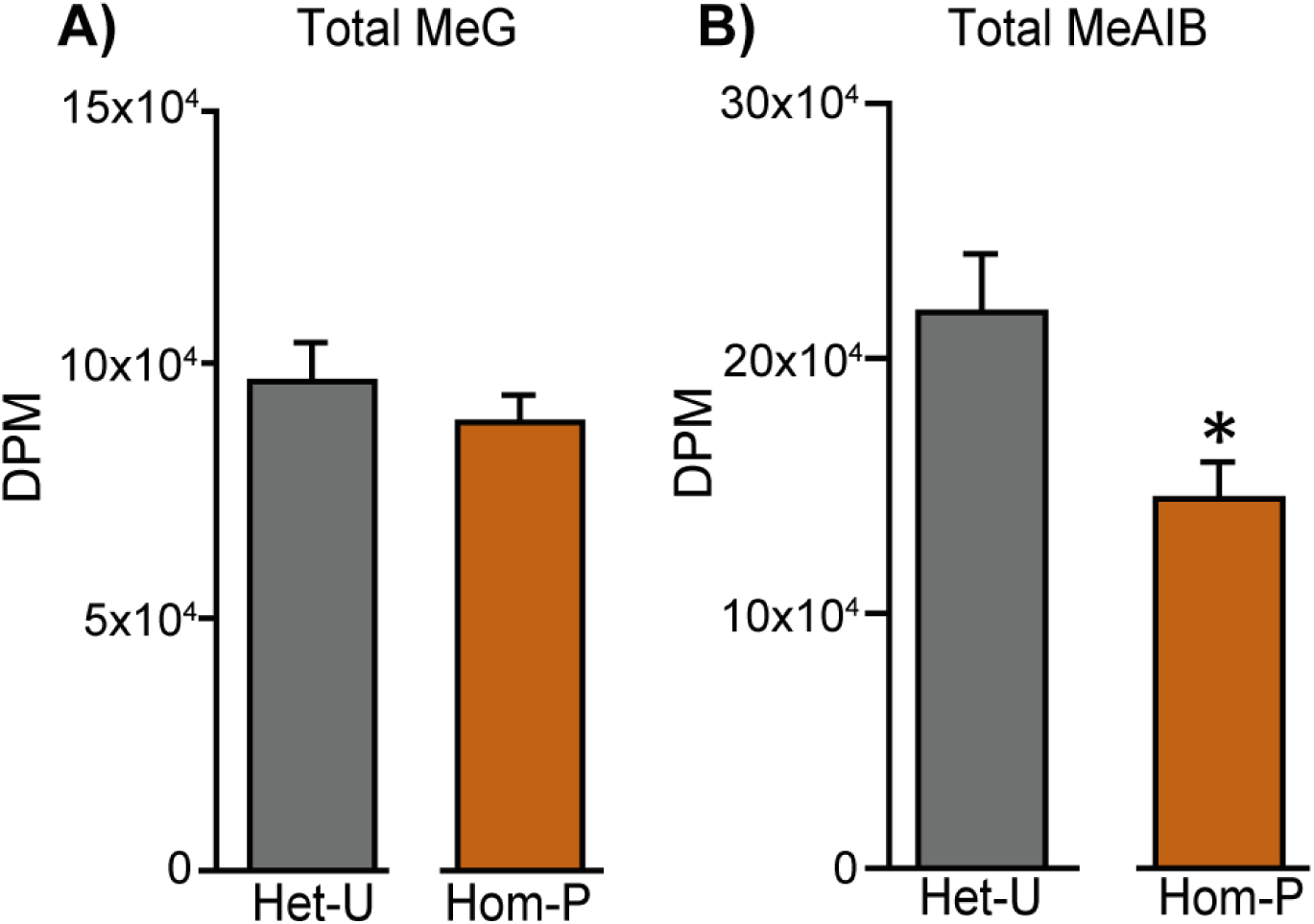
Total solute accumulation in Het-U versus Hom-P. MeG and MeAIB accumulation on day 19 of pregnancy in Het-U control values (n=16) and Hom-P (n=16). *P < 0.05 *versus* Het-U, unpaired t test

**Fig. S5.**
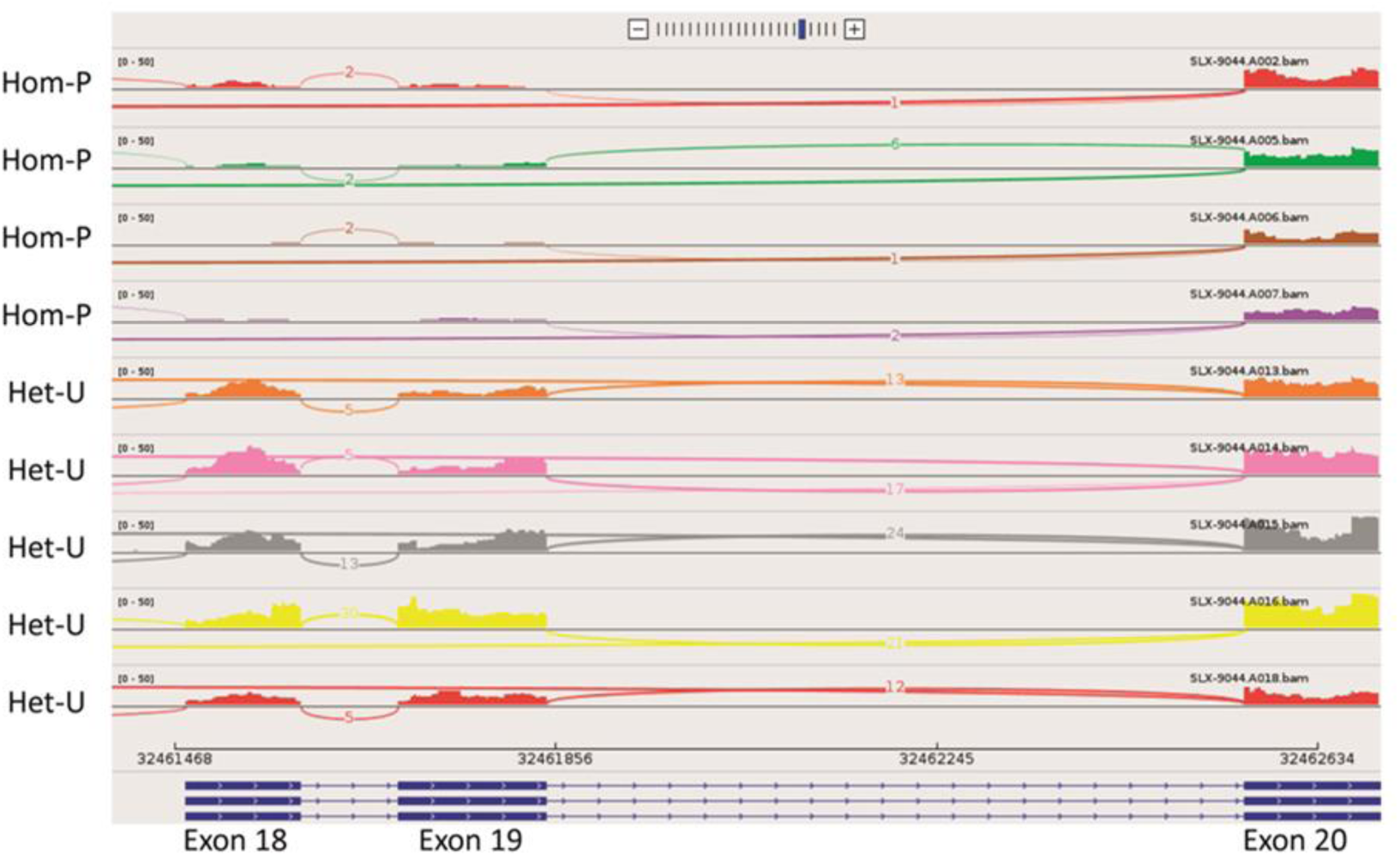
Sashimi plot showing RNA-seq reads for exons 18-20 of *Pik3ca* for the 4 Hom-P and 5 Het-U placentas analysed on day 19 of pregnancy.

**Fig. S6.**
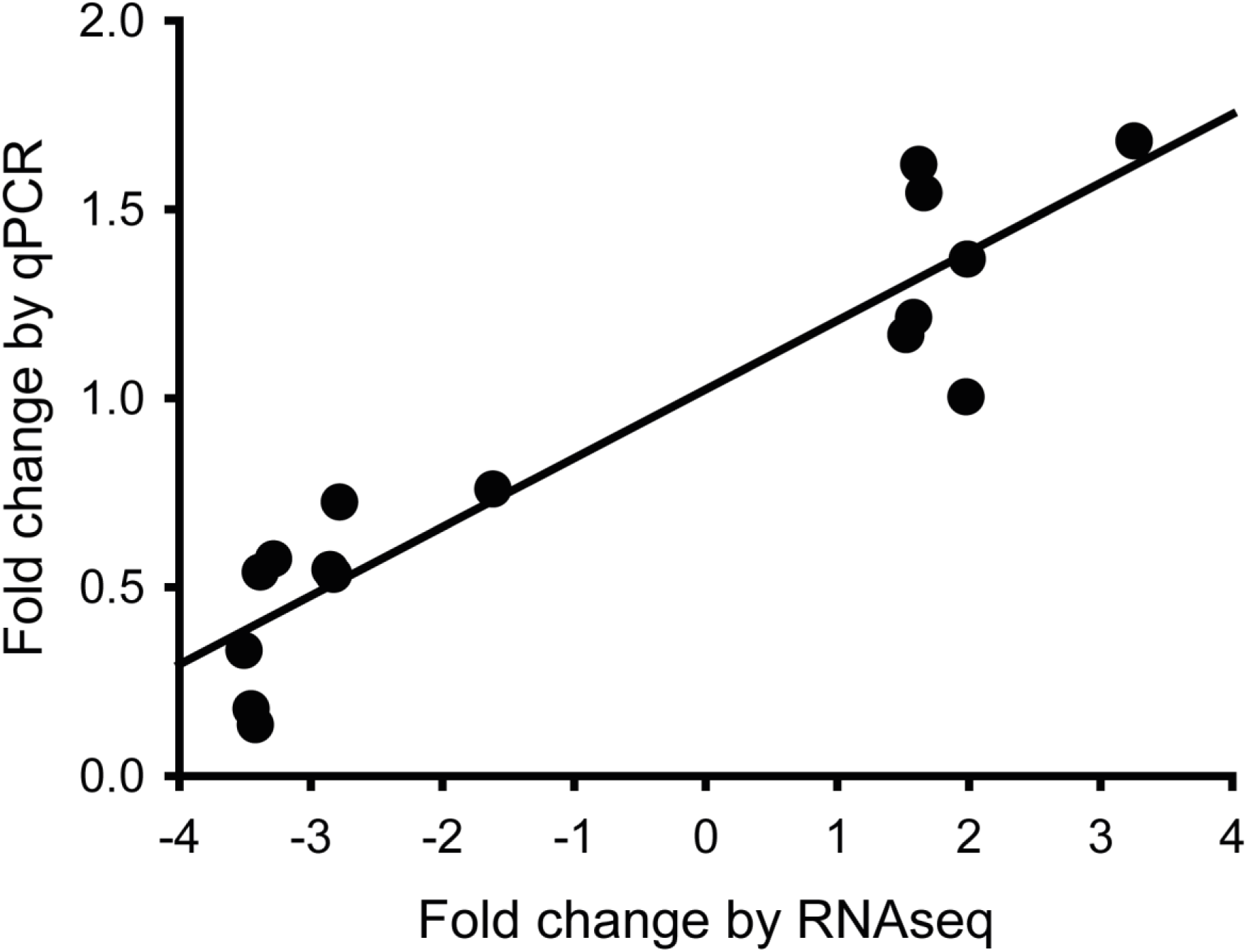
Validation of RNA-seq data. Validation performed by qRT-PCR for 16 genes in n=7 Het-U and Hom-P placentas on day 19 of pregnancy.

**Fig. S7.**
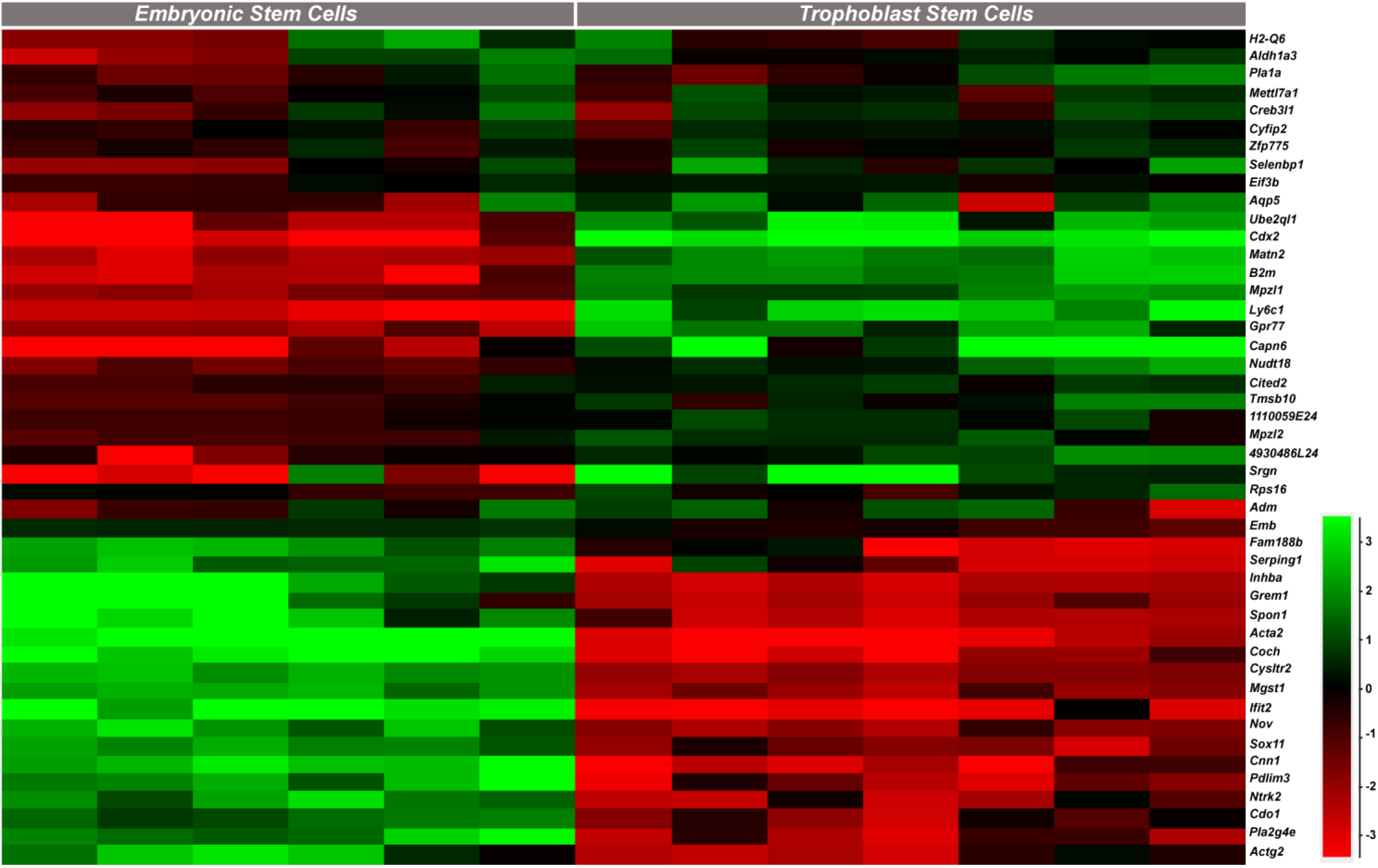
Expression profiles of the genes identified as dysregulated in Hom-P versus Het-U placentas in trophoblast stem cells (TS cells) and embryonic stem cells (ES cells), as determined by analysis of existing RNA-seq datasets (Chrysanthou et al., 2018; Latos et al., 2015).

**Table S1.** Summary of the mouse strains and experimental crosses used in the study. * Note for *Cyp19*Cre mutants (Het-P and Hom-P) the frequency is actually ∼half than stated, due to mosaic activity of this Cre line.

| Paternal genotype | Maternal genotype | Conceptus genotypes (for experiments) | Frequency* | Used for |
| --- | --- | --- | --- | --- |
| *Meox2Cre* (het) | *Pik3ca*<sup>Fl/Fl</sup> | *Pik3ca*<sup>Fl/+</sup> = WT<br>*Meox2Cre*; *Pik3ca*<sup>Fl/+</sup> = Het-F | 1:2 | Day 19 collection for fetal weight and placental transport phenotype |
| *Pik3ca* null (het);<br>*Meox2Cre* (het) | *Pik3ca*<sup>Fl/Fl</sup> | *Pik3ca*<sup>Fl/+</sup> = WT<br>*Meox2Cre*; *Pik3ca*<sup>Fl/+</sup> = Het-F<br>*Pik3ca* null het and *Pik3ca*<sup>Fl/+</sup> = Het-U<br>*Meox2Cre*; *Pik3ca*<sup>Fl/+</sup> and *Pik3ca* null het = Hom-F | 1:4 | D10-13 collection for conceptus viability |
| *Pik3ca*<sup>Fl/Fl</sup> | *Pik3ca* null (het);<br>*Cyp19Cre* (het) | *Pik3ca*<sup>Fl/+</sup> = WT<br>*Cyp19Cre*; *Pik3ca*<sup>Fl/+</sup> = Het-P<br>*Pik3ca* null het and *Pik3ca*<sup>Fl/+</sup> = Het-U<br>*Cyp19Cre*; *Pik3ca*<sup>Fl/+</sup> and *Pik3ca* null het = Hom-P | 1:4 | Day 19 collection for conceptus viability, fetal weight and placental transport phenotype |

**Table S2.** Deleting the remaining p110α from the trophoblast in Hom-P does not affect fetal viability at day 19 of pregnancy. Frequency of viable fetuses in a litter are displayed, with data from n=15 litters. Offspring genotypes were determined by conventional PCR, and in the case of *Cyp19*Cre mutants, additionally by qRT-PCR to identify those with a sufficient level of *Pik3ca* deletion (frequency is in parentheses). When the cut off for *Pik3ca* deletion in the placenta using qRT-PCR was applied (<65% for Het-P and <30% for Hom-P), the frequency of *Cyp19*Cre mutants was ∼50% less.

|  | WT | Het-P | Het-U | Hom-P |
| --- | --- | --- | --- | --- |
| Frequency by DNA genotyping (by qRT-PCR) | 26% | 23% (14%) | 19% | 32% (16%) |

**Table S3.** The effect of deleting the remaining p110α from the trophoblast in Hom-P on feto-placental growth relative to WT and Het-P. Hom-P * *versus* WT or † *versus* Het-P. *P < 0.05 and ***P < 0.001, †P < 0.05 and †††P < 0.001, unpaired t test. Conceptus weights are from n≥15, Lz and Jz volume from n≥6 and Lz morphology from n≥4 per genotype on day 19 of pregnancy. Data are presented as means ± SEM.

|  | WT | Het-P | Hom-P |
| --- | --- | --- | --- |
| Fetus (mg) | 1155±20 | 1152±25 | 994±22***††† |
| Placenta (mg) | 93±3 | 101±5 | 89±3† |
| Labyrinthine zone (mm<sup>3</sup>) | 47±2 | 46±4 | 36±2*** |
| Junctional zone (mm<sup>3</sup>) | 33±5 | 34±9 | 30±2 |
| Maternal blood spaces (mm<sup>3</sup>) | 14±2 | 9±1 | 11±1† |
| Trophoblast (mm<sup>3</sup>) | 28±2 | 34±2 | 22±2††† |
| Fetal capillaries (mm<sup>3</sup>) | 5±1 | 3±1 | 4±1*† |
| Barrier thickness (μm) | 3.5±0.1 | 3.7±0.1 | 3.9±0.1* |
| Surface area (mm<sup>2</sup>) | 15±1 | 9±1 | 10±1*** |

**Table S4.**
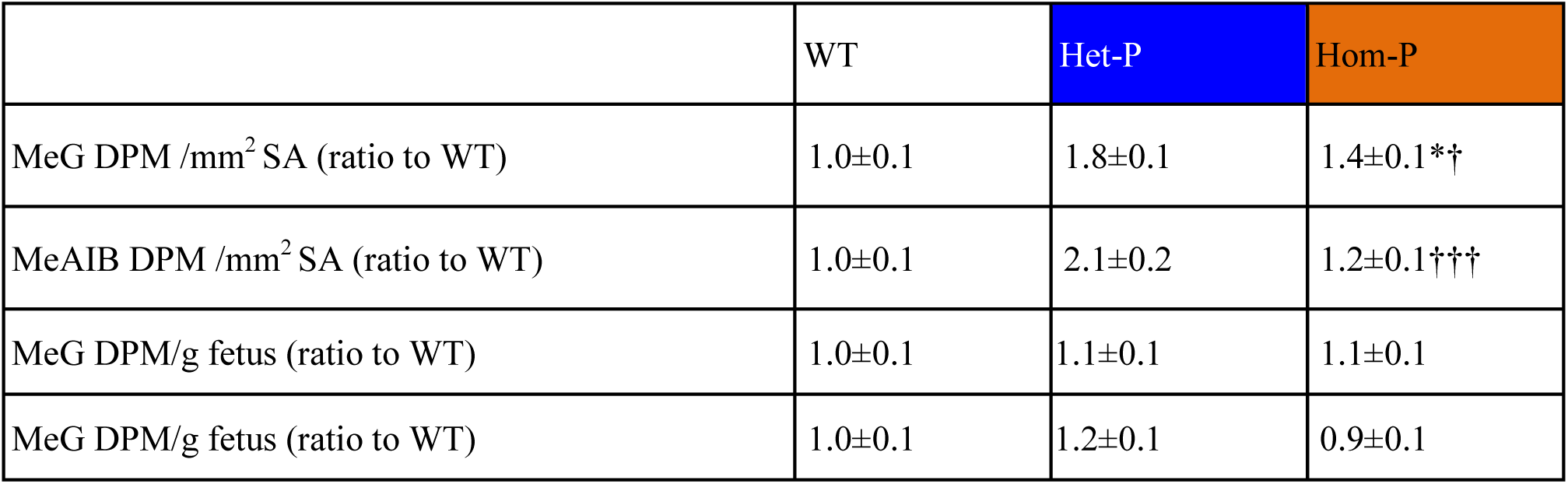
The effect of deleting the remaining p110α from trophoblast in Hom-P on placental transport capacity relative to WT and Het-P. Placental transport of ^3^H-methyl-D glucose (MeG) and 14C-amino isobutyric acid (MeAIB) relative to surface area available or to fetal weight on day 19 of pregnancy is shown as a ratio of WT values. Hom-P * *versus* WT or † *versus* Het-P. *P < 0.05, †P < 0.05 and †††P < 0.001, unpaired t test. Data are from n≥15 and presented as means ± SEM.

**Table S5.** List of differentially expressed genes obtained by RNA-seq comparing 5 Het-U and 4 Hom-P placentas on day 19 of pregnancy.

EXCEL DOCUMENT

**Table S6. Functional predictions for the candidate genes enriched in TS and ES cells**.

EXCEL DOCUMENT

**Table S7.**
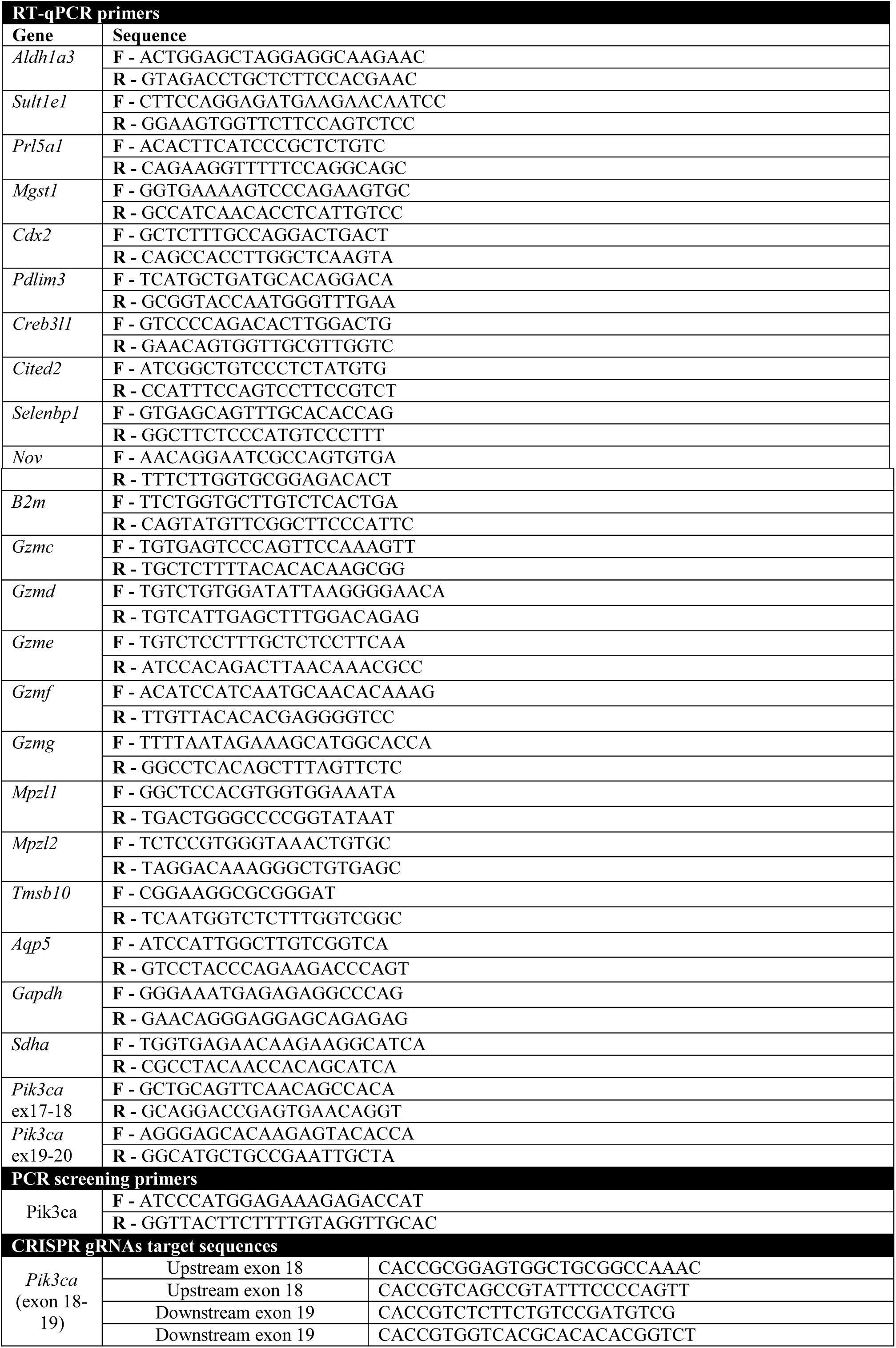
Primer sequence for RT-qPCR, PCR screening and CRISPR gRNAs target.

## References and notes

Adamson, S.L., Lu, Y., Whiteley, K.J., Holmyard, D., Hemberger, M., Pfarrer, C., and Cross, J.C. (2002). Interactions between trophoblast cells and the maternal and fetal circulation in the mouse placenta. Dev Biol 250, 358–373.

Angiolini, E., Coan, P.M., Sandovici, I., Iwajomo, O.H., Peck, G., Burton, G.J., Sibley, C.P., Reik, W., Fowden, A.L., and Constancia, M. (2011). Developmental adaptations to increased fetal nutrient demand in mouse genetic models of Igf2-mediated overgrowth. FASEB 25, 1737–1745.

Baschat, A.A., Galan, H.L., Ross, M.G., and Gabbe, S.G. (2007). Intrauterine growth restriction. In Obstetrics Normal and Problem Pregnancies, S.G. Gabbe, J.R. Niebyl, and J.L. Simpson, eds. (Churchill Livingstone Elsevier), pp. 771–814.

Baschat, A.A., and Hecher, K. (2004). Fetal growth restriction due to placental disease. Semin Perinatol 28, 67–80.

Bi, L., Okabe, I., Bernard, D.J., Wynshaw-Boris, A., and Nussbaum, R.L. (1999). Proliferative defect and embryonic lethality in mice homozygous for a deletion in the p110alpha subunit of phosphoinositide 3-kinase. Journal of Biological Chemistry 274, 10963–10968.

Blanco, B., Herrero-Sanchez, M.C., Rodriguez-Serrano, C., Sanchez-Barba, M., and Del Canizo, M.C. (2015). Profound blockade of T cell activation requires concomitant inhibition of different class I PI3K isoforms. Immunol Res 62, 175–188.

Chrysanthou, S., Senner, C.E., Woods, L., Fineberg, E., Okkenhaug, H., Burge, S., Perez-Garcia, V., and Hemberger, M. (2018). A Critical Role of TET1/2 Proteins in Cell-Cycle Progression of Trophoblast Stem Cells. Stem Cell Reports 10, 1355–1368.

Clark, D.A., Ding, J.W., Yu, G., Levy, G.A., and Gorczynski, R.M. (2001). Fgl2 prothrombinase expression in mouse trophoblast and decidua triggers abortion but may be countered by OX-2. Mol Hum Reprod 7, 185–194.

Coan, P.M., Angiolini, E., Sandovici, I., Burton, G.J., Constancia, M., and Fowden, A.L. (2008). Adaptations in placental nutrient transfer capacity to meet fetal growth demands depend on placental size in mice. Journal of Physiology 586, 4567–4576.

Coan, P.M., Ferguson-Smith, A.C., and Burton, G.J. (2004). Developmental dynamics of the definitive mouse placenta assessed by stereology. Biol Reprod 70, 1806–1813.

Constancia, M., Angiolini, E., Sandovici, I., Smith, P., Smith, R., Kelsey, G., Dean, W., Ferguson-Smith, A., Sibley, C.P., Reik, W., et al. (2005). Adaptation of nutrient supply to fetal demand in the mouse involves interaction between the Igf2 gene and placental transporter systems. Proc Natl Acad Sci USA 102, 19219–19224.

Constancia, M., Hemberger, M., Hughes, J., Dean, W., Ferguson-Smith, A., Fundele, R., Stewart, F., Kelsey, G., Fowden, A., Sibley, C., et al. (2002). Placental-specific IGF-II is a major modulator of placental and fetal growth. Nature 417, 945–948.

Cramer, S., Beveridge, M., Kilberg, M., and Novak, D. (2002). Physiological importance of system A-mediated amino acid transport to rat fetal development. Am J Physiol Cell Physiol 282, C153–160.

Diaz, L.E., Chuan, Y.-C., Lewitt, M., Fernandez-Perez, L., Carrasco-Rodriguez, S., Sanchez-Gomez, M., and Flores-Morales, A. (2007). IGF-II regulates metastatic properties of choriocarcinoma cells through the activation of the insulin receptor. Mol Hum Reprod 13, 567–576.

Dilworth, M.R., Kusinski, L.C., Cowley, E., Ward, B.S., Husain, S.M., Constancia, M., Sibley, C.P., and Glazier, J.D. (2010). Placental-specific Igf2 knockout mice exhibit hypocalcemia and adaptive changes in placental calcium transport. Proc Natl Acad Sci USA 107, 3894–3899.

Efeyan, A., Comb, W.C., and Sabatini, D.M. (2015). Nutrient-sensing mechanisms and pathways. Nature 517, 302–310.

Efimova, O.V., and Kelley, T.W. (2009). Induction of granzyme B expression in T-cell receptor/CD28-stimulated human regulatory T cells is suppressed by inhibitors of the PI3K-mTOR pathway. BMC Immunol 10, 59.

Engelman, J.A., Luo, J., and Cantley, L.C. (2006). The evolution of phosphatidylinositol 3-kinases as regulators of growth and metabolism. Nat Rev Genet 7, 606–619.

Espinoza, J., Sebire, N.J., McAuliffe, F., Krampl, E., and Nicolaides, K.H. (2001). Placental villus morphology in relation to maternal hypoxia at high altitude. Placenta 22, 606–608.

Forbes, K., Westwood, M., Baker, P.N., and Aplin, J.D. (2008). Insulin-like growth factor I and II regulate the life cycle of trophoblast in the developing human placenta. Am J Physiol Cell Physiol 294, C1313–1322.

Foukas, L.C., Claret, M., Pearce, W., Okkenhaug, K., Meek, S., Peskett, E., Sancho, S., Smith, A.J.H., Withers, D.J., and Vanhaesebroeck, B. (2006). Critical role for the p110[alpha] phosphoinositide-3-OH kinase in growth and metabolic regulation. Nature 441, 366–370.

Frevert, E.U., and Kahn, B.B. (1997). Differential effects of constitutively active phosphatidylinositol 3-kinase on glucose transport, glycogen synthase activity, and DNA synthesis in 3T3-L1 adipocytes. Molecular and cellular biology 17, 190–198.

Gellhaus, A., Schmidt, M., Dunk, C., Lye, S.J., Kimmig, R., and Winterhager, E. (2006). Decreased expression of the angiogenic regulators CYR61 (CCN1) and NOV (CCN3) in human placenta is associated with pre-eclampsia. Mol Hum Reprod 12, 389–399.

Georgiades, P., Cox, B., Gertsenstein, M., Chawengsaksophak, K., and Rossant, J. (2007). Trophoblast-specific gene manipulation using lentivirus-based vectors. Biotechniques 42, 317–318, 320, 322-315.

Glazier, J.D., Cetin, I., Perugino, G., Ronzoni, S., Grey, A.M., Mahendran, D., Marconi, A.M., Pardi, G., and Sibley, C.P. (1997). Association between the activity of the system A amino acid transporter in the microvillous plasma membrane of the human placenta and severity of fetal compromise in intrauterine growth restriction. Pediatr Res 42, 514–519.

Godfrey, K.M., Matthews, N., Glazier, J., Jackson, A., Wilman, C., and Sibley, C.P. (1998). Neutral amino acid uptake by the microvillous plasma membrane of the human placenta is inversely related to fetal size at birth in normal pregnancy. The Journal of clinical endocrinology and metabolism 83, 3320–3326.

Graupera, M., Guillermet-Guibert, J., Foukas, L.C., Phng, L.-K., Cain, R.J., Salpekar, A., Pearce, W., Meek, S., Millan, J., Cutillas, P.R., et al. (2008). Angiogenesis selectively requires the p110-a isoform of PI3K to control endothelial cell migration. Nature 453, 662–666.

Higgins, J.S., Vaughan, O.R., de Liger, E.F., Fowden, A.L., and Sferruzzi-Perri, A.N. (2015). Placental phenotype and resource allocation to fetal growth are modified by the timing and degree of hypoxia during mouse pregnancy. J Physiol 594, 1341–1356.

Huang da, W., Sherman, B.T., and Lempicki, R.A. (2009). Systematic and integrative analysis of large gene lists using DAVID bioinformatics resources. Nat Protoc 4, 44–57.

Jackson, M.R., Mayhew, T.M., and Haas, J.D. (1988). On the factors which contribute to thinning of the villous membrane in human placentae at high altitude. I. Thinning and regional variation in thickness of trophoblast. Placenta 9, 1–8.

Jansson, N., Pettersson, J., Haafiz, A., Ericsson, A., Palmberg, I., Tranberg, M., Ganapathy, V., Powell, T.L., and Jansson, T. (2006). Down-regulation of placental transport of amino acids precede the development of intrauterine growth restriction in rats fed a low protein diet. Journal of Physiology 576, 935–946.

Jansson, T., and Powell, T.L. (2006). IFPA 2005 Award in Placentology Lecture. Human placental transport in altered fetal growth: does the placenta function as a nutrient sensor? -- a review. Placenta 27 Suppl A, S91–97.

Jean, S., and Kiger, A.A. (2014). Classes of phosphoinositide 3-kinases at a glance. J Cell Sci 127, 923–928.

Jiang, K., Zhong, B., Gilvary, D.L., Corliss, B.C., Hong-Geller, E., Wei, S., and Djeu, J.Y. (2000). Pivotal role of phosphoinositide-3 kinase in regulation of cytotoxicity in natural killer cells. Nat Immunol 1, 419–425.

Katagiri, H., Asano, T., Ishihara, H., Inukai, K., Shibasaki, Y., Kikuchi, M., Yazaki, Y., and Oka, Y. (1996). Overexpression of catalytic subunit p110alpha of phosphatidylinositol 3-kinase increases glucose transport activity with translocation of glucose transporters in 3T3-L1 adipocytes. J Biol Chem 271, 16987–16990.

Kent, L., Konno, T., and Soares, M. (2010). Phosphatidylinositol 3 kinase modulation of trophoblast cell differentiation. BMC developmental biology 10, 97.

Kent, L.N., Ohboshi, S., and Soares, M.J. (2012). Akt1 and insulin-like growth factor 2 (Igf2) regulate placentation and fetal/postnatal development. Int J Dev Biol 56, 255–261.

Knight, Z.A., Gonzalez, B., Feldman, M.E., Zunder, E.R., Goldenberg, D.D., Williams, O., Loewith, R., Stokoe, D., Balla, A., Toth, B., et al. (2006). A pharmacological map of the PI3-K family defines a role for p110[alpha] in insulin signaling. Cell 125, 733–747.

Kriplani, N., Hermida, M.A., Brown, E.R., and Leslie, N.R. (2015). Class I PI 3-kinases: Function and evolution. Adv Biol Regul 59, 53–64.

Latos, P.A., Sienerth, A.R., Murray, A., Senner, C.E., Muto, M., Ikawa, M., Oxley, D., Burge, S., Cox, B.J., and Hemberger, M. (2015). Elf5-centered transcription factor hub controls trophoblast stem cell self-renewal and differentiation through stoichiometry-sensitive shifts in target gene networks. Genes & development 29, 2435–2448.

Lelievre, E., Bourbon, P.-M., Duan, L.-J., Nussbaum, R.L., and Fong, G.-H. (2005). Deficiency in the p110{alpha} subunit of PI3K results in diminished Tie2 expression and Tie2-/-like vascular defects in mice. Blood 105, 3935– 3938.

Limbourg, F.P., Takeshita, K., Radtke, F., Bronson, R.T., Chin, M.T., and Liao, J.K. (2005). Essential role of endothelial Notch1 in angiogenesis. Circulation 111, 1826–1832.

Lu, J., Zhang, S., Nakano, H., Simmons, D.G., Wang, S., Kong, S., Wang, Q., Shen, L., Tu, Z., Wang, W., et al. (2013). A positive feedback loop involving Gcm1 and Fzd5 directs chorionic branching morphogenesis in the placenta. PLoS Biol 11, e1001536.

Miinea, C.P., Sano, H., Kane, S., Sano, E., Fukuda, M., Peranen, J., Lane, W.S., and Lienhard, G.E. (2005). AS160, the Akt substrate regulating GLUT4 translocation, has a functional Rab GTPase-activating protein domain. Biochem J 391, 87–93.

Moore, L., David Young, D., McCullough, R., Droma, T., and Zamudio, S. (2001). Tibetan protection from intrauterine growth restriction (IUGR) and reproductive loss at high altitude. American Journal of Human Biology 13, 635–644.

Moore, T., and Dveksler, G.S. (2014). Pregnancy-specific glycoproteins: complex gene families regulating maternal-fetal interactions. Int J Dev Biol 58, 273–280.

Moreau, J.L., Artap, S.T., Shi, H., Chapman, G., Leone, G., Sparrow, D.B., and Dunwoodie, S.L. (2014). Cited2 is required in trophoblasts for correct placental capillary patterning. Dev Biol 392, 62–79.

Mounayar, M., Kefaloyianni, E., Smith, B., Solhjou, Z., Maarouf, O.H., Azzi, J., Chabtini, L., Fiorina, P., Kraus, M., Briddell, R., et al. (2015). PI3kalpha and STAT1 Interplay Regulates Human Mesenchymal Stem Cell Immune Polarization. Stem Cells 33, 1892–1901.

Okada, Y., Ueshin, Y., Isotani, A., Saito-Fujita, T., Nakashima, H., Kimura, K., Mizoguchi, A., Oh-Hora, M., Mori, Y., Ogata, M., et al. (2007). Complementation of placental defects and embryonic lethality by trophoblast-specific lentiviral gene transfer. Nat Biotechnol 25, 233–237.

Pantham, P., Rosario, F.J., Weintraub, S.T., Nathanielsz, P.W., Powell, T.L., Li, C., and Jansson, T. (2016). Down-regulation of placental transport of amino acids precedes the development of intrauterine growth restriction in maternal nutrient restricted baboons. Biol Reprod 95, 98.

Plaks, V., Berkovitz, E., Vandoorne, K., Berkutzki, T., Damari, G.M., Haffner, R., Dekel, N., Hemmings, B.A., Neeman, M., and Harmelin, A. (2011). Survival and size are differentially regulated by placental and fetal PKBalpha/AKT1 in mice. Biol Reprod 84, 537–545.

Rosario, F.J., Kanai, Y., Powell, T.L., and Jansson, T. (2012). Mammalian target of rapamycin signalling modulates amino acid uptake by regulating transporter cell surface abundance in primary human trophoblast cells. Journal of Physiology 591, 609–625.

Sahay, A.S., Sundrani, D.P., Wagh, G.N., Mehendale, S.S., and Joshi, S.R. (2015). Neurotrophin levels in different regions of the placenta and their association with birth outcome and blood pressure. Placenta 36, 938–943.

Sandovici, I., Hoelle, K., Angiolini, E., and Constancia, M. (2012). Placental adaptations to the maternal-fetal environment: implications for fetal growth and developmental programming. Reprod Biomed Online 25, 68–89.

Schwenk, F., Baron, U., and Rajewsky, K. (1995). A cre-transgenic mouse strain for the ubiquitous deletion of loxP-flanked gene segments including deletion in germ cells. Nucleic Acids Res 23, 5080–5081.

Sferruzzi-Perri, A.N., and Camm, E.J. (2016). The programming power of the placenta. Front Physiol 7:33.

Sferruzzi-Perri, A.N., Lopez-Tello, J., Fowden, A.L., and Constancia, M. (2016). Maternal and fetal genomes interplay through phosphoinositol 3-kinase(PI3K)-p110α signalling to modify placental resource allocation. Proc Natl Acad Sci USA 113(40), 11255–11260.

Sferruzzi-Perri, A.N., Sandovici, I., Constancia, M., and Fowden, A.L. (2017). Placental phenotype and the insulin-like growth factors: resource allocation to fetal growth. Journal of Physiology 595, 5057–5093.

Sferruzzi-Perri, A.N., Vaughan, O.R., Coan, P.M., Suciu, M.C., Darbyshire, R., Constancia, M., Burton, G.J., and Fowden, A.L. (2011). Placental-specific Igf2 deficiency alters developmental adaptations to undernutrition in mice. Endocrinology 152, 3202–3212.

Sferruzzi-Perri, A.N., Vaughan, O.R., Forhead, A.J., and Fowden, A.L. (2013a). Hormonal and nutritional drivers of intrauterine growth. Curr Opin Clin Nutr Metab Care 16, 298–309.

Sferruzzi-Perri, A.N., Vaughan, O.R., Haro, M., Cooper, W.N., Musial, B., Charalambous, M., Pestana, D., Ayyar, S., Ferguson-Smith, A.C., Burton, G.J., et al. (2013b). An obesogenic diet during mouse pregnancy modifies maternal nutrient partitioning and the fetal growth trajectory. FASEB 27, 3928–3937.

Sheu, M.L., Shen, C.C., Jheng, J.R., and Chiang, C.K. (2017). Activation of PI3K in response to high glucose leads to regulation of SOCS-3 and STAT1/3 signals and induction of glomerular mesangial extracellular matrix formation. Oncotarget 8, 16925–16938.

Sibley, C.P., Coan, P.M., Ferguson-Smith, A.C., Dean, W., Hughes, J., Smith, P., Reik, W., Burton, G.J., Fowden, A.L., and Constancia, M. (2004). Placental-specific insulin-like growth factor 2 (Igf2) regulates the diffusional exchange characteristics of the mouse placenta. Proc Natl Acad Sci USA 101, 8204–8208.

Soares, M.J., Konno, T., and Alam, S.M.K. (2007). The prolactin family: effectors of pregnancy-dependent adaptations. Trends in Endocrinology & Metabolism 18, 114–121.

Sopasakis, V.R., Liu, P., Suzuki, R., Kondo, T., Winnay, J., Tran, T.T., Asano, T., Smyth, G., Sajan, M.P., Farese, R.V., et al. (2010). Specific roles of the p110alpha isoform of phosphatidylinsositol 3-kinase in hepatic insulin signaling and metabolic regulation. Cell Metab 11, 220–230.

Soriano, P. (1999). Generalized lacZ expression with the ROSA26 Cre reporter strain. Nat Genet 21, 70–71.

Strumpf, D., Mao, C.-A., Yamanaka, Y., Ralston, A., Chawengsaksophak, K., Beck, F., and Rossant, J. (2005). Cdx2 is required for correct cell fate specification and differentiation of trophectoderm in the mouse blastocyst. Development 132, 2093–2102.

Supek, F., Bosnjak, M., Skunca, N., and Smuc, T. (2011). REVIGO summarizes and visualizes long lists of gene ontology terms. PLoS One 6, e21800.

Tallquist, M.D., and Soriano, P. (2000). Epiblast-restricted Cre expression in MORE mice: a tool to distinguish embryonic vs. extra-embryonic gene function. Genesis 26, 113–115.

Tang, N., Mack, F., Haase, V.H., Simon, M.C., and Johnson, R.S. (2006). pVHL function is essential for endothelial extracellular matrix deposition. Molecular and cellular biology 26, 2519–2530.

Terragni, J., Nayak, G., Banerjee, S., Medrano, J.L., Graham, J.R., Brennan, J.F., Sepulveda, S., and Cooper, G.M. (2011). The E-box binding factors Max/Mnt, MITF, and USF1 act coordinately with FoxO to regulate expression of proapoptotic and cell cycle control genes by phosphatidylinositol 3-kinase/Akt/glycogen synthase kinase 3 signaling. J Biol Chem 286, 36215–36227.

Than, N.G., Romero, R., Tarca, A.L., Kekesi, K.A., Xu, Y., Xu, Z., Juhasz, K., Bhatti, G., Leavitt, R.J., Gelencser, Z., et al. (2018). Integrated Systems Biology Approach Identifies Novel Maternal and Placental Pathways of Preeclampsia. Front Immunol 9, 1661.

Todros, T., Marzioni, D., Lorenzi, T., Piccoli, E., Capparuccia, L., Perugini, V., Cardaropoli, S., Romagnoli, R., Gesuita, R., Rolfo, A., et al. (2007). Evidence for a role of TGF-beta1 in the expression and regulation of alpha-SMA in fetal growth restricted placentae. Placenta 28, 1123–1132.

Vanhaesebroeck, B., Guillermet-Guibert, J., Graupera, M., and Bilanges, B. (2010). The emerging mechanisms of isoform-specific PI3K signalling. Nat Rev Mol Cell Biol 11, 329–341.

Vaughan, O.R., Sferruzzi-Perri, A.N., Coan, P.M., and Fowden, A.L. (2013). Adaptations in placental phenotype depend on route and timing of maternal dexamethasone administration in mice. Biol Reprod 89, 1–12.

Wang, J., Knight, Z.A., Fiedler, D., Williams, O., Shokat, K.M., and Pearce, D. (2008). Activity of the p110-{alpha} subunit of phosphatidylinositol-3-kinase is required for activation of epithelial sodium transport. Am J Physiol Renal Physiol 295, F843–850.

Wenzel, P.L., and Leone, G. (2007). Expression of Cre recombinase in early diploid trophoblast cells of the mouse placenta. Genesis 45, 129–134.

Withington, S.L., Scott, A.N., Saunders, D.N., Lopes Floro, K., Preis, J.I., Michalicek, J., Maclean, K., Sparrow, D.B., Barbera, J.P., and Dunwoodie, S.L. (2006). Loss of Cited2 affects trophoblast formation and vascularization of the mouse placenta. Dev Biol 294, 67–82.

Wyrwoll, C.S., Seckl, J.R., and Holmes, M.C. (2009). Altered placental function of 11{beta}-hydroxysteroid dehydrogenase 2 knockout mice. Endocrinology 150, 1287–1293.

Xu, X.-y., Zhang, Z., Su, W.-h., Zhang, Y., Feng, C., Zhao, H.-m., Zong, Z.-h., Cui, C., and Yu, B.-z. (2009). Involvement of the p110α isoform of PI3K in early development of mouse embryos. Molecular reproduction and development 76, 389–398.

Yang, Z.-Z., Tschopp, O., Hemmings-Mieszczak, M., Feng, J., Brodbeck, D., Perentes, E., and Hemmings, B.A. (2003). Protein kinase B{alpha}/Akt1 regulates placental development and fetal growth. J Biol Chem 278, 32124– 32131.

Zhang, Q., Chen, L., Zhao, Z., Wu, Y., Zhong, J., Wen, G., Cao, R., Zu, X., and Liu, J. (2018). HMGA1 Mediated High-Glucose-Induced Vascular Smooth Muscle Cell Proliferation in Diabetes Mellitus: Association Between PI3K/Akt Signaling and HMGA1 Expression. DNA Cell Biol 37, 389–397.

